# Deep Proteome Profiling Enabled Functional Annotation and Data-Independent Quantification of Proline Hydroxylation Targets

**DOI:** 10.1101/2022.01.12.474993

**Authors:** Yao Gong, Gaurav Behera, Luke Erber, Ang Luo, Yue Chen

## Abstract

Proline hydroxylation (Hyp) regulates protein structure, stability and protein-protein interaction and is widely involved in diverse metabolic and physiological pathways in cells and diseases. To reveal functional features of the proline hydroxylation proteome, we integrated various data sources for deep proteome profiling of proline hydroxylation proteome in human and developed HypDB (https://www.HypDB.site), an annotated database and web server for proline hydroxylation proteome. HypDB provides site-specific evidence of modification based on extensive LC-MS analysis and literature mining with 15319 non-redundant Hyp sites and 8226 sites with high confidence on human proteins. Annotation analysis revealed significant enrichment of proline hydroxylation on key functional domains and tissue-specific distribution of Hyp abundance across 26 types of human organs and fluids and 6 cell lines. The network connectivity analysis further revealed a critical role of proline hydroxylation in mediating protein-protein interactions. Moreover, the spectral library generated by HypDB enabled data-independent analysis (DIA) of clinical tissues and the identification of novel Hyp biomarkers in lung cancer and kidney cancer. Taken together, our integrated analysis of human proteome with publicly accessible HypDB revealed functional diversity of Hyp substrates and provides a quantitative data source to characterize proline hydroxylation in pathways and diseases.

## 1. Introduction

Proline hydroxylation (Hyp), first discovered in 1902, is an important protein posttranslational modification pathway in cellular physiology and metabolism [1]. Simply adding a hydroxyl group to the imino side chain of proline residue, the modification is found to be evolutionarily conserved from bacteria to humans. In mammalian cells, proline hydroxylation is largely mediated through the enzymatic activities of two major families of prolyl hydroxylases – collagen prolyl 4-hydroxylases (P4HAs) [2] and hypoxia induced factor (HIF) prolyl hydroxylase domain proteins (PHDs) [3], while there are no known enzymes capable of removing proline hydroxylation discovered yet. Since the activity of prolyl hydroxylases depends on the cellular collaboration of multiple co-factors including oxygen, iron as well as several metabolites such as alpha-ketoglutarate, succinate and ascorbate, proline hydroxylation pathway is an important metabolic-sensing mechanism in the cells and tissues.

The most well-characterized Hyp targets are collagen proteins and HIFα family of transcription factors. Proline hydroxylation on collagens mediated by prolyl-4-hydroxylases (P4Hs) is critical to maintain the triple-helical structure of the collagen polymer and enable the proper protein folding after translation. Indeed, adding an electronegative oxygen on proline 4R position promotes the trans conformation and stabilizes the secondary structure of collagen [1a]. Inhibition of collagen proline hydroxylation destabilizes the collagen and prevent its export from the ER, therefore inducing cell stress and death [4]. HIFα transcription factors are essential to mediate hypoxia-response in mammalian cells [5]. Proline hydroxylation of HIFα proteins mediated by prolyl hydroxylase domain proteins (PHDs) under normoxia condition is recognized by pVHL in the Cullin 2 E3 ligase complex, which leads to rapid ubiquitination and degradation of HIFα proteins [6]. Hypoxia condition inhibits HIFα proline hydroxylation and degradation, enabling the transcriptional activation of over 100 hypoxia-responsing genes [7]. In the past two decades, numerous studies driven by advances in mass spectrometry-based proteomics technology have reported the identification and characterization of diverse new Hyp targets and the important roles of the modification in physiological functions [8]. Proline hydroxylation has been well known to affect protein homeostasis and the classic example is PHD-HIF-pVHL regulatory axis. The similar mechanism also regulates the turnover of diverse key transcriptional, metabolic and signaling proteins, including β2AR, NDRG3, ACC2, EPOR, G9a and SFMBT1 etc. [9]. In addition to pVHL-mediated protein degradation, proline hydroxylation also regulates substrate degradation by affecting its interaction with deubiquitinases. For example, the hydroxylation of Foxo3a promotes substrate degradation by inhibiting the interaction with deubiquinase Usp9x, and hydroxylation of p53 enhances its interaction with deubiquitinases Usp7/Usp10 to prevent its rapid degradation [10]. P4H-mediated proline hydroxylation has also been known to regulate the stability of diverse substrates including AGO2 and Carabin [11]. In addition to protein degradation, proline hydroxylation can also affect protein-protein interaction to regulate signaling and transcriptional activities. For example, PKM2 hydroxylation promotes its binding with HIF1A for transcriptional activation, proline hydroxylation of AKT enhances the interaction with pVHL to inhibit the kinase activity of AKT, and PHD1-mediated hydroxylation of Rpb1 is necessary for its translocation and phosphorylation [12]. More recently, TBK1 hydroxylation was identified and found to induce pVHL and phosphatase binding, which decreases its phosphorylation and enzyme activity, while the loss of pVHL hyperactivates TBK1 and promotes tumor development in clear cell renal cell carcinoma (ccRCC) [8d, 13].

Despite of these advances, there is a lack of integrated and annotated knowledgebase dedicated for proline hydroxylation, which underappreciates the functional diversity and physiological significance of this evolutionarily conserved metabolic-sensing PTM pathway. To fill the knowledge gap, we developed a publicly accessible proline hydroxylation database, HypDB (http://www.HypDB.site) (**Figure S1**). The development of the HypDB provides three main features – first, a classification-based algorithm for confident identification of proline hydroxylation substrates, second, integrated resources based on exhaustive manual literature mining, large-scale LCMS analysis, curated public database, and third, a collection of a large spectral library for LCMS-based site-specific identification from a variety of cell lines and tissues. Furthermore, stoichiometry-based quantification of Hyp sites allows quantitative comparison of site abundance across various proteins and tissues, and the extensively annotated proline hydroxylation proteome enables deep bioinformatic analysis including network connectivity, structural domain enrichment and tissue-specific distribution study. The online database system allows the community-driven submission of LCMS dataset to be included in HypDB annotation and the direct export of precursor and fragmentation with spectral library that enables the development of targeted quantitative proteomics and data-independent analysis workflow. We hope that the HypDB will provide critical insights into the functional diversity and network of the proline hydroxylation proteome and aid in further mechanistic studies on the physiological roles of the metabolic-sensing PTM pathway in cells and diseases.

## 2. Results

### 2.1 Database Construction and Analysis Workflow

To construct a bioinformatic resource for metabolic-sensing proline hydroxylation targets, we developed HypDB, a MySQL-based relational database on a public-accessible web server (**Figure 1 and S2**). It was constructed based on three main resources to comprehensively annotate human proline hydroxylation proteome (**Figure 1**). First, manual curation of literature through Pubmed (searching term: “proline hydroxylation” and time limit between 2000 and 2021) was performed by two independent curators, which yielded 1287 research journal articles. Site identification was extracted from each journal article and its corresponding protein was mapped to Uniprot protein ID if possible. Manual curation of the research articles focused on the sites that were biochemically investigated with multiple evidence including mass spectrometry, mutagenesis, Western blotting as well as *in vitro* or *in vivo* enzymatic assays. Analyzed Hyp site identifications were then matched against the existing data in the database to reduce redundancy. Second, the database included extensive LCMS-based direct evidence of proline hydroxylation site identifications based on the integrated analysis of over 100 LCMS datasets of various human cell lines and tissues (See Experimental Methods). The datasets were either downloaded from publicly accessible server or produced in-house. Each dataset was analyzed through a standardized workflow using Maxquant search engine and the Hyp site identifications were filtered and imported into the HypDB with a bioinformatic analysis pipeline specified in details below. Our collection of MS-based evidence of Hyp identifications from cell lines and tissues likely revealed a significant portion of Hyp sites that can be potentially identified by deep proteomic analysis as evidenced by our observation that the rate of unique Hyp site addition from each dataset decreased significantly despite the increased collection of datasets in the database (**Figure S2B**). Third, the HypDB also integrated Hyp identification annotated in the public Uniprot database. To better clarification, the database records whether the site was uniquely reported by Uniprot database or also identified by large-scale LCMS analysis.

**Figure 1.**
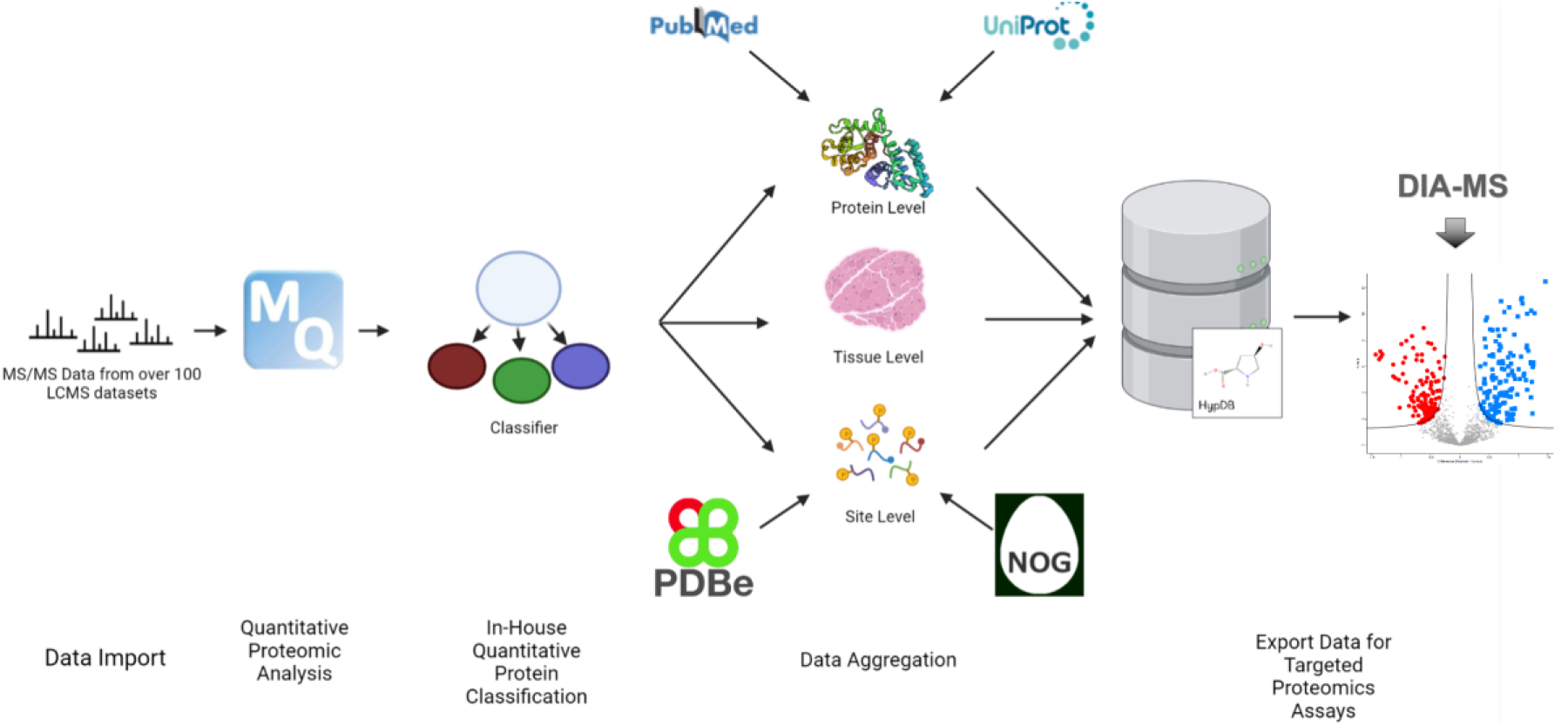
Workflow of establishing HypDB database and webserver. The process included deep proteome profiling analysis of human tissues and cell lines, manual literature mining and integration with Uniprot data source. Classification-based algorithm was applied to extract confident identifications and site-specific bioinformatic analysis with stoichiometr-based quantification revealed the biochemical pathways involved with human Hyp proteome. MS-based Hyp library further enabled DIA-MS quantification of Hyp proteome in cells and tissues.

We implemented stringent criteria for data importing and classification from LCMS-based identifications. To import data into the HypDB, LCMS-based identification of Hyp site from database search analysis was first analyzed by a classification-based algorithm to determine the confidence of Hyp site identification and localization (**Figure 2A**). The classification was performed using the best scored MS/MS spectrum of a Hyp site in each dataset analysis. The algorithm classified Hyp identifications that can be exclusively localized to proline residue based on consecutive b-or y-ions as Class I sites. The algorithm classified the Hyp identifications that cannot be exclusively localized based on MS/MS spectrum analysis but can be distinguished from five common types of oxidation artifacts ( Methionine, Tryptophan, Tyrosine, Histidine, Phenylalanine) mainly induced during sample preparation as Class II sites. Other Hyp identifications that were reported by MaxQuant database search software (with 1% false-discovery rate at the site-level and a minimum Andromeda score of 40) were grouped as Class III sites. We further developed a site-localization score using the relative intensities of key fragment ion to index the level of confidence in site-localization with MS/MS spectrum analysis for Class I and Class II sites (**Experimental Methods**). Each dataset was analyzed by the classification algorithm separately and the best classification evidence for each Hyp site was selected and reported on the HypDB website to indicate the confidence of site localization. The classification-based algorithm provides the specificity and reliability required for an accurately annotated database while maintaining all possible identifications as searchable records.

**Figure 2.**
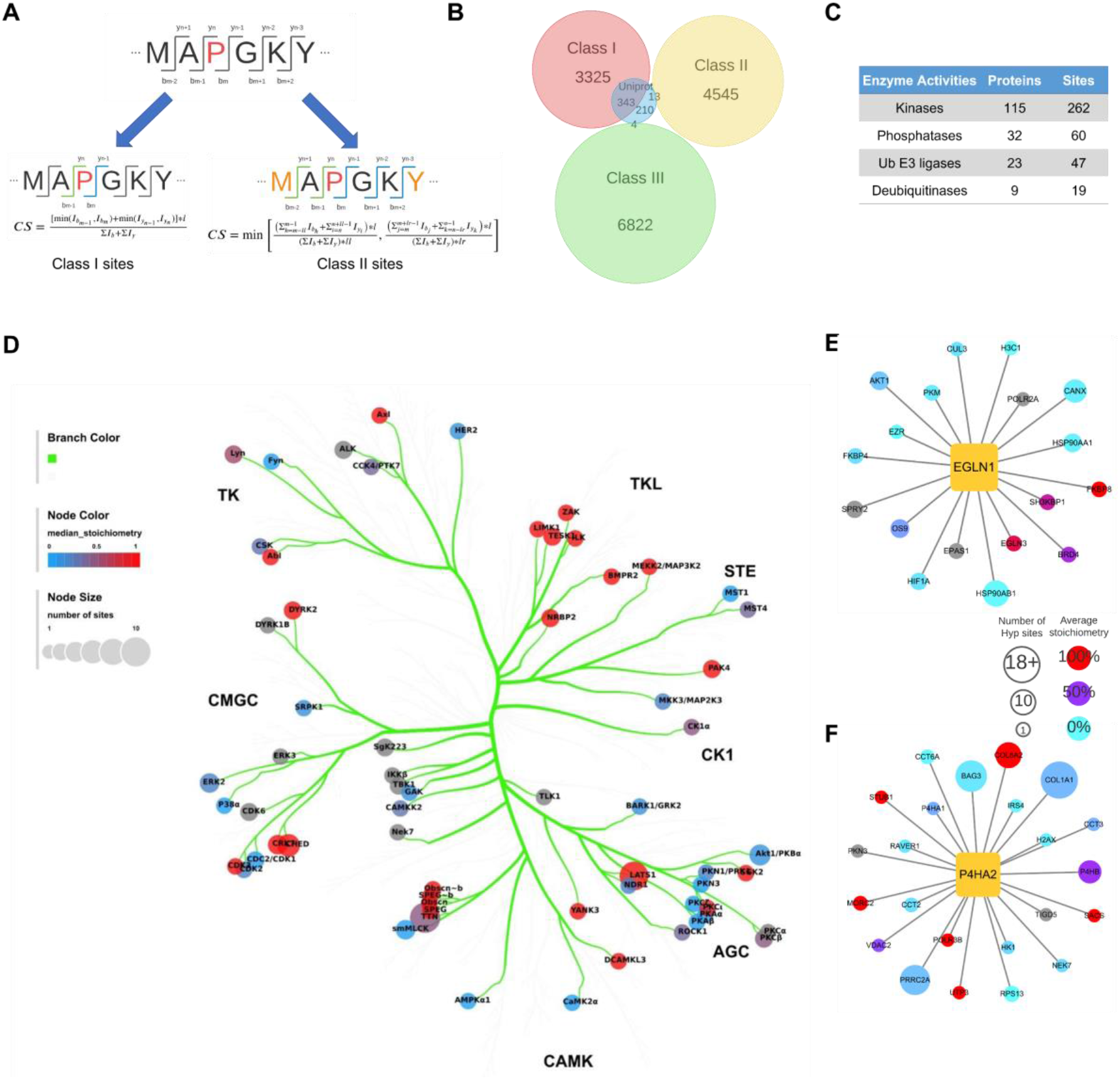
Substrate diversity of human Hyp proteome. A. Illustration of classification based algorithm to identify confident Hyp sites; B.Venn diagram of Class I, II, III Hyp sites identified from MS analysis and manually curated Uniprot sites; C. PTM regulatory enzymes identified as Hyp substrates; D. Kinase tree classification showing the distributions of kinases as Hyp substrates in different kinase families, including AGC (named after PKA, PKG, PKC families), CAMK (leaded by calcium/calmodulin-dependent protein kinases), CK1 (cell kinase 1), CMGC (named after CDKs, MAPK, GSK, CLK families), STE (Homologs of the yeast STE counterparts), TK (tyrosine kinases), and TKL (tyrosine kinase-like); E. Hydroxyproline proteins that interact with EGLN1; F. Hydroxyproline proteins that interact with P4HA2.

To evaluate the site-specific prevalence of proline hydroxylation, a stoichiometry-based quantification strategy was integrated into the analysis workflow using the previously established principles [8d, 14]. Briefly, the Hyp stoichiometry was calculated by dividing the summed intensities of the peptides with the Hyp site identification with the total intensities of the peptides containing the same proline site in the dataset. HypDB recorded all available site-specific Hyp stoichiometry analysis from various cell lines and tissues, which allowed site-specific quantitative analysis of modification abundance across cell and tissue types. The stoichiometry distribution of Class I and Class II sites were shown in **Figure S2C-D**, and the median stoichiometry of all stoichiometry measurements for any specific site was calculated and reported on the HypDB website.

To further explore the functional association of Hyp proteome, several bioinformatic annotation strategies were also integrated into the analysis workflow. These stand-alone workflows include evolutionary conservation analysis, solvent accessibility analysis and protein-protein interface analysis. Evolutionary conservation analysis compared the conservation of Hyp sites with other proline sites on the same protein and performed statistical test to determine if the Hyp site is more evolutionarily conserved than non-Hyp sites. Solvent accessibility analysis analyzed the sequence of the substrate protein with DSSP package and calculated the likelihood of solvent accessibility for each Hyp sites. Protein-protein interaction interface analysis extracted the domain interaction residues from the 3DID database based on PDB structure analysis and matched them against the Hyp site in the database to identify the Hyp site that is localized in the interface and more likely to interfere with protein-protein interaction.

All information above was integrated into several tables and linked through foreign keys as the schema in **Figure S2A**. Complete information on all Hyp sites was organized in two major tables including a redundant-site table (**Table S1**), which stored all Hyp sites identified in different tissues and cell lines including annotated MS/MS spectra, site-specific abundance and sample source information, and a nonredundant-site table (**Table S2**), which merged the LCMS-based evidence from different sources at the site-specific level and also integrated with the sites collected from Uniprot and manual curation of literatures.

### 2.2 Validation of the Hyp Site Classification Strategy

To validate our classification-based strategy for confidence Hyp site identification, we performed comparative analysis of Hyp site identifications from each class with manually curated Uniprot Hyp identifications. Our analysis showed that the Class I and II Hyp sites covered over 60 percent sites annotated in the Uniprot, while very few Uniprot annotated sites overlapped with the Class III sites, suggesting that our Hyp site localization and classification algorithm allowed the collection of highly confident Hyp identification and significantly improved the reliability of LCMS-based Hyp site analysis (**Figure 2B**). To further probe the current state of the proline hydroxylation proteome, we performed bioinformatic analysis for functional annotation based on highly confident Hyp site identifications in HypDB.

### 2.3 Mapping human proline hydroxylation proteome

HypDB currently collected 15319 non-redundant Hyp sites out of 60209 Hyp site records through large-scale deep proteomics analysis of different tissue, cell lines, manual curation of literatures and integration with Uniprot database. Among 15293 non-redundant Hyp sites, 3668 sites were categorized as Class I sites, 4558 sites were categorized as Class II sites, 6826 were categorized as Class III sites (**Figure 2B**). In addition, the database contained 57 sites from literature mining and 210 sites that were integrated from the Uniprot database. Excluding Class III proline hydroxylation sites, we identified a total of 115 kinases (262 sites), 32 phosphatases (60 sites), 23 E3 ligases (47 sites) and 9 deubiquitinases (19 sites) as Hyp substrates (**Figure 2C**). Statistical analysis showed a specific enrichment of kinases in proline hydroxylation proteome (p=0.0393), suggesting a potentially broad crosstalk between proline hydroxylation and kinase signaling pathways (**Figure 2D**). Comparing Hyp substrates with the interactome of prolyl hydroxylases in BioGRID [15], we identified 18 Hyp proteins with 52 sites that are known to interact with EGLN1/PHD2, 16 Hyp proteins with 33 sites that are known to interact with EGLN2/PHD1, 297 Hyp proteins with 636 sites that are known to interact with EGLN3/PHD3, 35 Hyp proteins with 98 sites that are known to interact with P4HA1, 24 Hyp proteins with 173 sites that are known to interact with P4HA2, and 19 Hyp proteins with 56 sites that are known to interact with P4HA3 (**Figure 2E-F**).

To determine if proline hydroxylation site is more accessible to solvent, we collected 3D structures of proteins from PDBe and UniProt and calculated the relative solvent accessibility (RSA) of each proline residual on proteins with hydroxyproline sites with the DSSP package [16]. To examine if there’s an RSA difference between Hyp sites and non-Hyp sites on protein with Hyp sites, we performed a two-tail t-test and found no significant difference in the distribution of solvent accessibility, suggesting that proline hydroxylation does not necessarily target solvent accessible proline residues (**Figure S3A**). To determine if proline hydroxylation targets proline sites that are more evolutionarily conserved, we performed evolutionary conservation analysis through extensive sequence alignment of protein orthologs across species based on EggNOG database [17] and statistically compare the conservation of Hyp sites with the conservation of all proline on the same protein. Our data showed that about 49% sites were evolutionarily conserved with statistical significance (p<0.05) (**Figure S3B**). To determine if Hyp could play a potential role in domain-domain interactions, we analyzed data of known domain-based interactions of three-dimensional protein structures from HypDB non-redundant site database (excluding Class III Hyp sites). We identified 168 unique Hyp sites that were located at the interface of the interaction. These data suggesting potential involvement of proline hydroxylation in directly regulating protein-protein interaction. For example, proline hydroxylation at position 14 on Superoxide dismutase (SOD1) will form a hydrogen-bonding with a neighboring chain Gln16 in a dimeric structure and potentially promote the stabilization of the dimer (**Figure S3C**).

### 2.3 Functional Features of Proline Hydroxylation Proteins

We performed GO enrichment tests and other functional annotations on proteins that contain Class I, II, literature or Uniprot sites (**Figure 3A**). Our analysis revealed that Hyp substrates are highly enriched in metabolic processes such as response to toxic substances (p<10^-24^) and organic cyclic compound catabolic process (p<10^-21^), mRNA splicing (p<10^-26^) and structural functions such as NABA collagens (p<10^-35^), supramolecular fiber organization (p<10^-50^), and cell morphogenesis involved in differentiation (p<10^-31^). To determine if proline hydroxylation proteome prefers to be involved in protein-protein interactions, we extracted a human protein interaction database from STRING with a cutoff score of 0.7 and then extracted all the interactions containing two Hyp proteins based on the STRING database. Based on these data, we performed network connectivity analysis by comparing the number of interactions of Hyp proteins with the distribution of the number of interactions from randomly selected human proteins with 10,000 times of repeats. Our data showed that Hyp substrates are significantly involved in the protein-protein network (p<0.0001) (**Figure 3B**). We further performed protein complex enrichment analysis using manually curated CORUM database, and our analysis showed that Hyp proteome is significantly enriched with many known protein complexes (**Figure S6**), such as TNF-alpha/NF-kappa B signaling complex 6 (**Figure S7A**), TLE1 corepressor complex (**Figure S7B**), DGCR8 multiprotein complex (**Figure S7C**), Nop56p-associated pre-rRNA complex (**Figure S7D**), and PA700-20S-PA28 complex (**Figure S7E**), suggesting that proline hydroxylation targets proteins in multiple pathways that affects signaling and gene expression. Using MCODE clustering analysis, we extracted significantly enriched clusters from Hyp proteome interaction network and these highly connected clusters of Hyp substrates suggested that proline hydroxylation targets important cellular activities including regulation of mRNA splicing, hypoxia response, and focal adhesion (**Figure 3C-E**).

**Figure 3.**
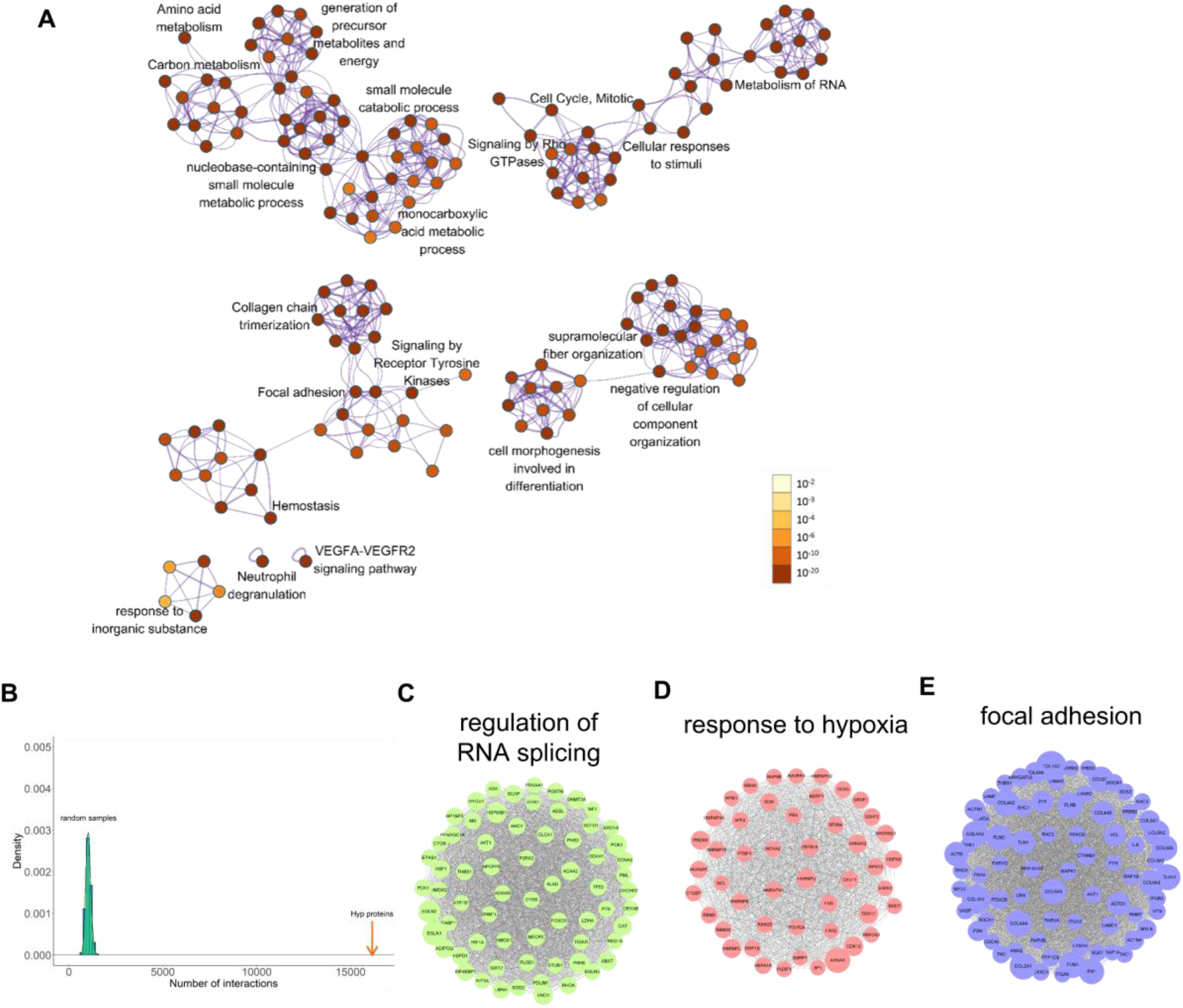
Gene enrichment and connectivity analysis of HypDB. A. Interaction network of top 20 enriched term clusters of HypDB; B. Bootstrapping-based analysis of hydroxyproline protein interactions comparing to a distribution of protein interactions from random samples with the same number of human proteins; C. Hydroxyproline proteins enriched in the regulation of RNA splicing; D. Hydroxyproline proteins enriched in the response to hypoxia; E. Hydroxyproline proteins enriched in focal adhesion.

### 2.4 Structural and Motif Features of Proline Hydroxylation Sites

We analyzed the local sequence context around Hyp sites (excluding Class III sites) using the MoMo software tool [18] As we expected, Hyp sites with PG motif and GPPG motif were highly enriched (p<10^-10^) which is characteristic for collagen protein families (**Figure 4A and S4A**). In addition to collagen, we identified 33 proteins with similar motif to collagen and these proteins may be potential substrates of prolyl-4-hydroxylases. Other than the collagen-like motif, we also identified CP motif (p < 10^-6^) (**Figure 4A**) and proteins containing CP motifs are highly enriched in focal adhesion (FDR<0.05). To remove the high background of sites with collagen-like Hyp motifs, we filtered out sites with local sequence contexts in PG motif. Our re-analysis identified that acidic amino acids were enriched at the +1 position to form PD motif (**Figure 4A**). PD motif containing proteins were highly enriched in metabolic pathways (FDR<0.05). As proline hydroxylation sites may have crosstalk with other protein. Our analysis revealed 2372 phosphorylation sites and 532 ubiquitination sites that have been identified very close to the Hyp sites (**Figure S4B**).

**Figure 4.**
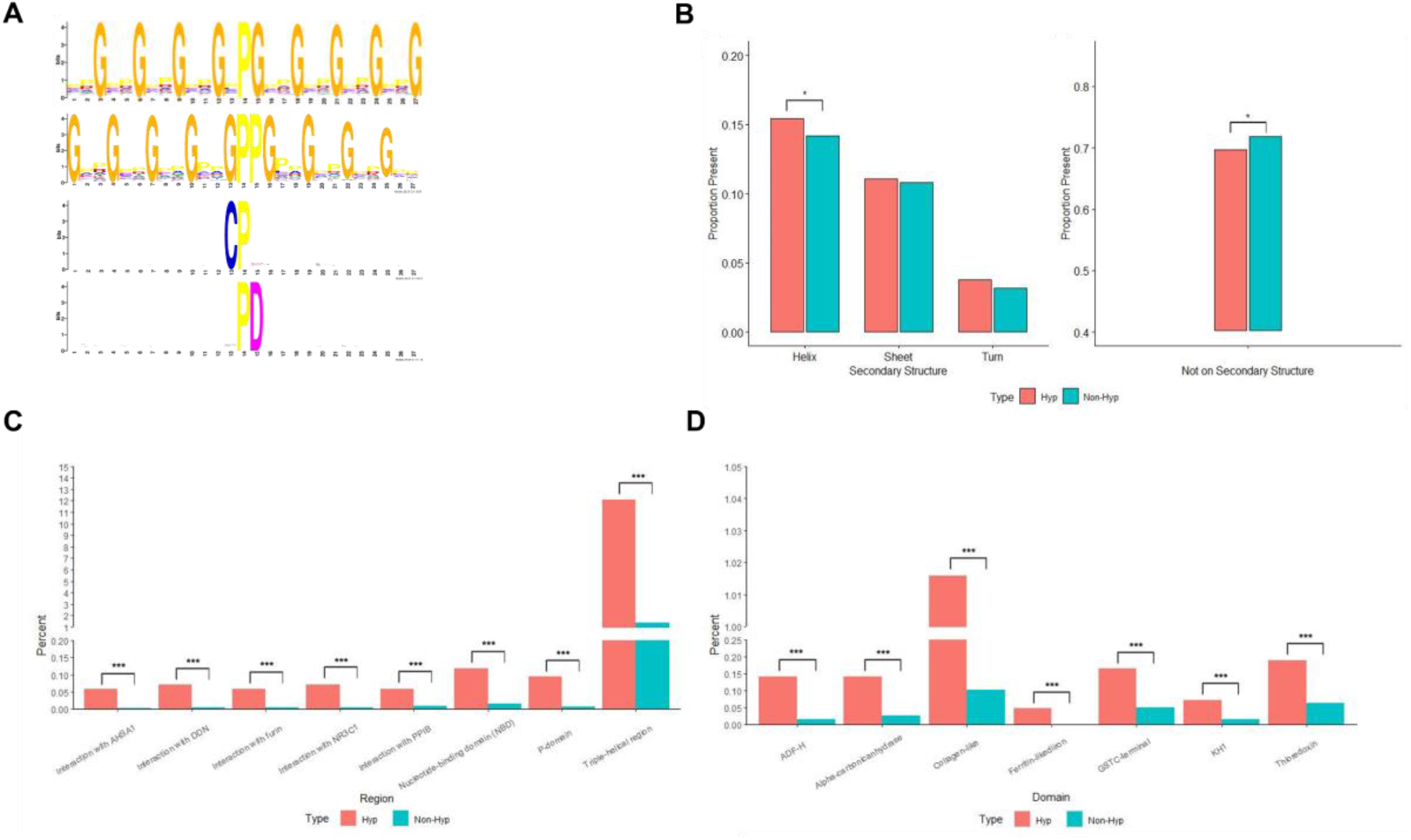
Motif and Protein Feature Analysis of HypDB. A. Consensus motifs for Hyp sites with top 3 motifs generated with all sites within HypDB, p < 10^-6^ and the bottom motif identified with Hyp sites after filtering out sequences with PG motif, p < 10^-6^; B. Proline residues located within secondary structures of proteins in HypDB. (* p<0.05); C. Proline residues located within regions of proteins in HypDB (*** p<0.001); D. Proline residues located within domains of proteins in HypDB (*** p<0.001)

To determine the structural features of proline hydroxylation sites, we extracted all Hyp proteins with known secondary structures. These proteins contain 2234 Hyp sites and 26012 Non-Hyp sites on sequences that have experimentally determined PDB structure. We then classified structure features into helix, sheet, coil and non-structure regions and performed statistical analysis to compare the secondary structure features of Hyp sites and non-Hyp sites. We found that proline hydroxylation indeed preferentially targets proline residues that are localized in helix secondary structure (p<0.05) (**Figure 4B** left panel). Accordingly, we observed a depletion of Hyp sites outside of a secondary structure feature (**Figure 4B** right panel).

As secondary structure may not fully represent functional structural features, we developed similar statistical analysis strategy to determine the site-specific enrichment of Hyp sites on functional domains or structural regions. In contrast to the traditional domain enrichment analysis using Pfam or Interpro for protein-level analysis, our strategy enabled site-specific enrichment analysis of domains or regions based on Uniprot annotation. Application of this strategy revealed diverse known and novel structural features that were highly enriched with proline hydroxylation, such as the triple-helical region, which is characteristic for collagen protein family (**Figure 4C**). In addition to the triple-helical region, our analysis revealed more than 10 functional regions and domains that were highly enriched with proline hydroxylation, including p-domain (p<10^-6^), NBD domain (p<10^-6^), thioredoxin domain (p<10^-3^) and ferritin-like domain (p<10^-6^) (**Figure 4C-D**). These data revealed previously unexpected role of proline hydroxylation targeting functional domains in diverse cellular pathways.

### 2.5 Site-specific Stoichiometric Quantification of Hyp Proteome

Comparing to relative quantification, stoichiometry analysis measures the prevalence and dynamics of the modification in a physiologically meaningful manner [8d, 19]. Our mass spectrometry-based deep proteome profiling enables site-specific quantification of proline hydroxylation stoichiometries across multiple tissues and cell lines. Our data showed that site-specific abundance of proline hydroxylation varies widely from below 1% to nearly 100% with an overall median stoichiometry of 7.66% (**Figure 5A**). Indeed, a bulk portion of the Hyp sites have either very low or very high stoichiometries. To investigate the functional differences between sites with different stoichiometry, we divided proteins into five quantiles based on average stoichiometry measurement for the same site across all cells and tissues (**Figure 5B**). The four cutoffs four cutoffs 5%, 30%, 70%, and 95% were selected so that each quantile contained a similar number of Hyp sites. We then performed GO enrichment and functional annotation on the five quantiles respectively and performed hierarchical clustering with Euclidean distance. Our data showed that proteins in immune response and neutrophil activation pathways are enriched with low to medium stoichiometry and proteins in cell adhesion and system development are enriched with medium to high stoichiometry (**Figure 5B**). We also saw a significant enrichment of proteins involved in chromatin assembly and RNA processing but the stoichiometry of hydroxylation on those proteins seems to be very low (**Figure 5B**). Combining site-specific functional feature annotation and stoichiometry analysis, we performed stoichiometry-based clustering of Hyp-targeted functional domains. Our data showed that ODD region which is known to regulate hydroxylation-mediated protein degradation of HIFα was enriched with medium stoichiometry and triple helical region on collagen, whose hydroxylation is required for its maturation, was enriched with high stoichiometry (**Figure 5C**). Furthermore, our analysis revealed stoichiometry-based enrichment of kinase domains at medium stoichiometry, GATA1 interaction domains at high stoichiometry, nucleotide binding domains at low to medium stoichiometry and histone binding domains at low stoichiometry (**Figure 5C**).

**Figure 5.**
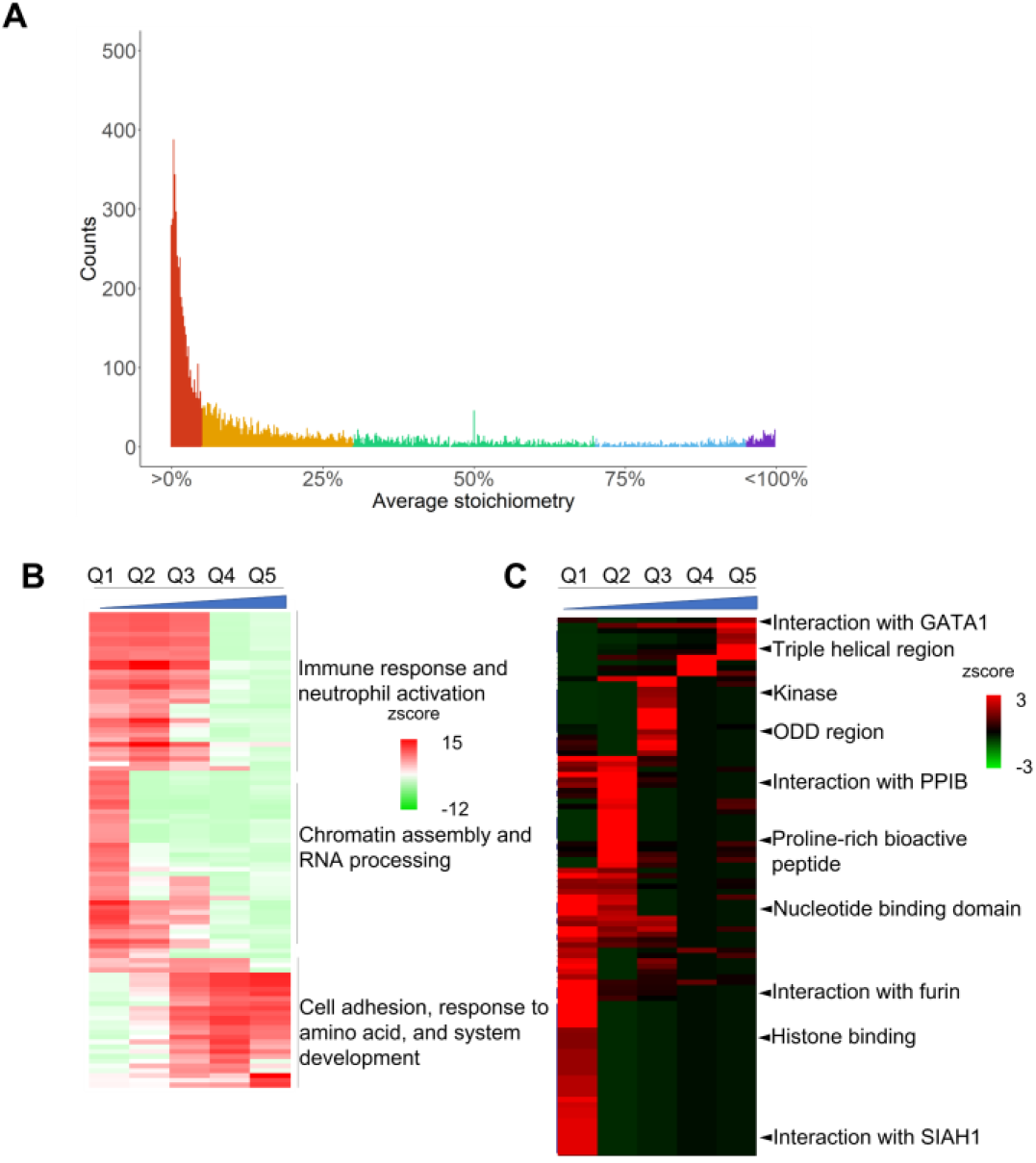
Enrichment analysis of different stoichiometry quantiles of HypDB. A. Stoichiometry distribution of hydroxyproline proteins that was divided into five quantiles – Q1, Q2, Q3, Q4, and Q5, from low to high stoichiometry with four cutoffs of 5%, 30%,70%, and 95% respectively; B. Heatmap representation of GO biological processes enrichment among five quantiles; C. Heatmap representation of domain and region enrichment among five quantiles.

### 2.6 Tissue-specific Distribution of Hyp Proteome

The collection of mass spectrometry-based identification of proline hydroxylation proteome enabled cross-tissue comparative analysis (**Table S3**). Indeed, at individual protein level, we observed a wide distribution of proline hydroxylation abundance for the same site and between different sites across different tissue (**Figure 6A and S5**). For example, Fibrillin-1 (FBN1) was identified with 22 Hyp sites of which 17 were Class I or II sites. Hyp1090 on EGF_CA repeat showed consistent high Hyp stoichiometry (71% to 96%) across four different tissues (testis, colon, heart, rectum), while Hyp1453 on another EGF_CA repeat showed varied Hyp stoichiometry (3% to 50.5%) across the same four tissues (testis, colon, heart and rectum) (**Figure 6A**). In another example, 6-phosphogluconate dehydrogenase (PGD) was identified with eight Hyp sites with half of them belonging to Class I or II sites. Hyp169 on the NAD binding domain showed relatively low stoichiometries in heart, liver and ovary (7.6% to 11.6%) but much higher stoichiometries in gut and B-cell (21.9% and 75.6%) (**Figure S5B**). We performed pathway enrichment analysis of proline hydroxylation and clustering of the enrichment across the tissues. Our data showed that proline hydroxylation proteome varied dramatically in terms of pathway and abundance among tissues (**Figure 6B-C**). For example, in lung, the Hyp proteome is mainly involved in collagen synthesis and tissue development, and it has relatively low portion of unique Hyp sites, but in liver, the Hyp proteome is heavily involved in diverse metabolic and translational processes with many liver specific proline hydroxylation targets (**Figure 6B-C**). Interestingly, clustering analysis showed that tissues sharing similar physiological functions tend to share similar proline hydroxylation profiles. Testis and ovary, for example, have similar enrichment of Hyp proteins related to DNA repair, gene silencing, and transcription regulation (**Figure 6D**). Urinary bladder and kidney, which also have similar enrichment patterns, contain Hyp proteins related to skeletal system morphogenesis and response to reactive oxygen species. One other tissue pair with similar Hyp proteins and enrichment results is spinal cord and frontal cortex; they are co-enriched in synapse organization and neuron projection development (**Figure 6D**).

**Figure 6.**
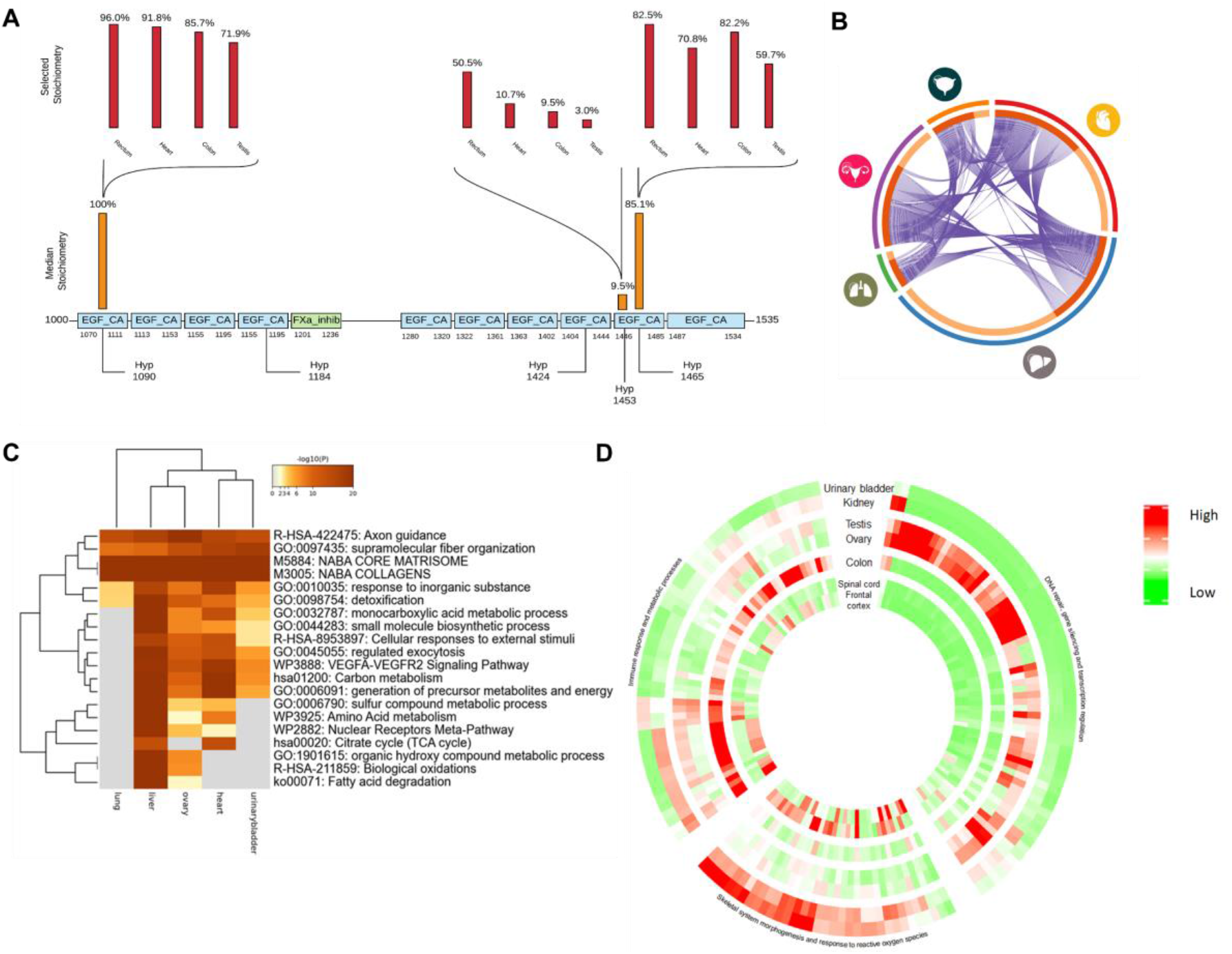
Proline hydroxylation proteome distributions in different tissues. A. An example showing varying stoichiometries of Hyp sites across different types of tissue for Fibrillin-1 (FBN1) with protein domains labeled in colored boxes; B. Correlation plot of Hyp proteins in five different tissues: heart, liver, lung, ovary, and urinary bladder with the size of arc shows relative number and the purple curved lines showing overlap proteins; C. Heatmap of Top 20 enrichment terms of hydroxyproline proteins found in five tissues; D. GO biological process enrichment heatmap of hydroxyproline proteins across six tissues.

### 2.7 Data-Independent Acquisition (DIA) Analysis of Hyp targets with HypDB-generated Spectral Library

Data-independent acquisition has been developed in the past ten years as a powerful strategy for reliable and efficient quantification of proteins and PTM sites [20]. Our extensive collection of the MS-based evidence for human proline hydroxylation sites provided an ideal resource to establish a data-independent acquisition workflow for global, site specific quantification of proline hydroxylation targets in cells and tissues. To this end, our web server has integrated functions for the direct export of annotated MS/MS identification of Hyp sites for selected proteins, cell line, tissue or at a proteome scale. The Export function provided two options – exporting the peptide precursor m/z only or exporting formatted MS/MS spectra. The former option can generate target m/z list that can be used as an inclusion list for targeted quantification of Hyp sites on selected proteins or sites. The latter option can directly generate spectral library used for data-independent acquisition analysis. Using the Export function, the current HypDB allowed the generation of a comprehensive Hyp spectral library in the NIST Mass Search format (msp) consisting of 6346 precursor ions, 5414 peptides, representing 8231 sites from 3222 proteins. To demonstrate the applicability of our resource in DIA analysis workflow, we analyzed two recently published large-scale DIA analysis datasets [20d, 20e]. Both datasets applied DIA analysis to quantify protein dynamics in the multiple replicates of paired normal and tumor samples.

The study by Kitata et al analyzed global protein profiles of lung cancer with five pairs of tumor and normal tissues in triplicate analysis for a total of thirty DIA-based LCMS runs [20d]. As a routine procedure in DIA analysis, we first performed database searching of DDA data in the dataset. Then, using this spectral library, we performed DIA analysis of all tumor and normal tissues with replicates. The analysis quantified 1422 Hyp sites from Kitata et al study (1% FDR). Next, we applied HypDB-generated spectral library and repeated the DIA analysis. Our result showed that using HypDB-generated spectral library led to a more than doubled the total number of Hyp sites quantified from DIA datasets using DDA-based spectral library with 3016 Hyp sites while covering more than 80% of the Hyp sites identified by the two DIA analysis, suggesting that the application of HypDB-generated spectral library was sufficient to cover majority of the Hyp identifications and significantly increased the sensitivity of Hyp proteome coverage (**Figure 7A**). DIA analysis with combined library generated by both HypDB and DDA identified 3651 Hyp sites and 1249 Hyp proteins (1% FDR). To determine the reproducibility of the quantification, we calculated the distribution of the percentage of coefficient variance (%CV) for DIA analysis of Hyp sites. Our data showed that %CV varied between 2% and 15% with a median value around 5% (**Figure 7B**), similar to the %CV distribution observed in the DIA analysis of proteins and phosphoproteins [20d]. Given the high reproducibility of the quantification, we filtered the Hyp sites with global 1% q-value cutoff (2283 sites) and performed hierachichal clustering analysis of Hyp sites quantified with normalized intensity in tumor and normal lung tissues (**Figure 7C**). Our data clearly showed that site-specific Hyp quantification was sufficient to cluster and distinguish tumor vs normal tissue. To identify significantly up-or down-regulated Hyp sites in tumor tissues, we performed two-samples t-test and analyzed the data in the volcano plot (**Figure 7D**). The analysis allowed us to identify 142 Hyp sites that were significantly upregulated and 179 Hyp sites that were significantly downregulated in tumor tissue (5% permutation-based FDR) (**Table S4**).. The dynamically regulated Hyp sites showed strong characteristics that were distinct between tumor and normal tissue. Interestingly, we observed subtype-dependent proline hydroxylation dynamics on collagen proteins. Collegen subtype IV, VI showed significantly downregulated proline hydroxylation level across multiple sites in tumor samples while collegen subtype X showed significantly increased proline hydroxylation (**Figure 7D**). Since proline hydroxylation promotes the structural stability of collagens, such changes likely indicated a signficantly increase in stability for Collagen X and decrease in stability for Collagen IV and VI in lung cancer tissue compared to the normal tissue. Our finding agreed well with a very recent publication indicating a pro-metastatic role of upregulated Collagen X in lung cancer progression [21]. In addition, we also identified significant upregulation of proline hydroxylation on glycolysis enzymes pyruvate kinase (PKM) and enolase (ENO1), heat-shock protein HSPA5 as well as autophagy protein Parkin (PARK7) (**Figure 7D**). P4HB, a member of the collagen prolyl 4-hydroxylase enzyme, also showed significant increase in proline hydroxylation (**Figure 7D**), likely due to increased prolyl 4-hydroxylase activity in lung cancer [22]. To reveal the functional significance of upregulated or downregulated Hyp substrates, we performed pathway and annotation enrichment analysis. Our data showed the pathways related to homeostatic processes, proteoglycan metabolism, cell-cell adhesion, regulation of stress-induced MAPK cascade and membrane hyperpolarization were significantly enriched among upregulated Hyp substrates (**Figure 7E**), while pathways related with collagen complexes and metabolism, negative regulation of angiogenesis, regulation of cell apoptosis, response to stimulus and cell morphogenesis were significantly enriched among down-regulated Hyp substrates (**Figure 7F**).

**Figure 7.**
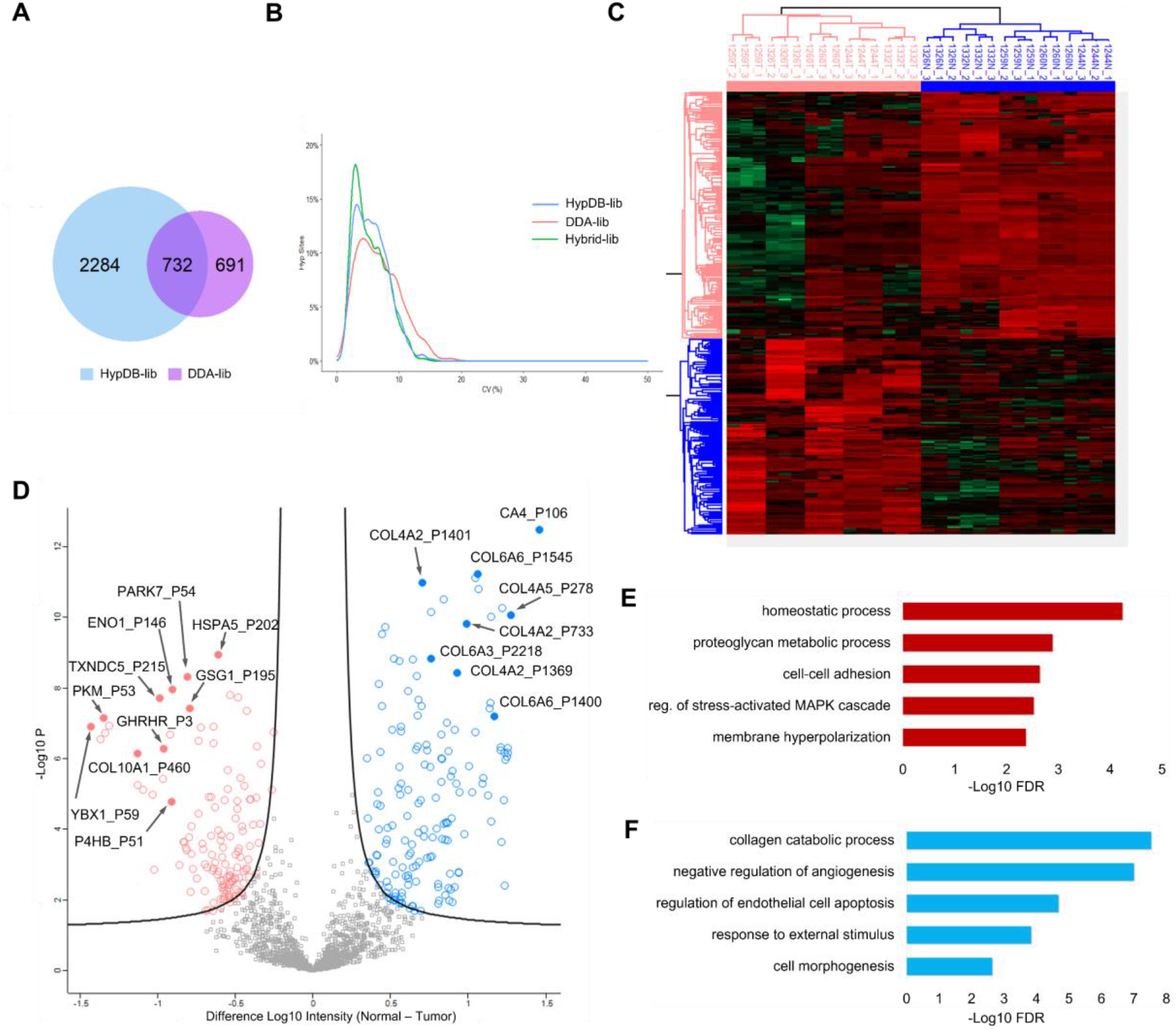
Analysis of Hyp proteome in lung cancer with DIA analysis. A. Venn diagram of DIA identifiation of Hyp sites using HypDB-generated library and the library generated by the Data-dependent acquisition (DDA) of the study; B. Distribution of %CV for Hyp sites quantified with HypDB-generated library, DDA-generated library or Hybrid librar combined with both sources; Hierachical clustering (C) and volcano plot (D) of significantly up or down-regulated Hyp sites in normal and tumor tissues; Significantly enriched GO biological processes among upregulated (E) and downregulated (F) proteins in tumor.

In another study, Guo et al applied DIA analysis to quantitatively profile kidney cancer proteome and the dataset consisted of 18 normal tissue analysis and 18 tumor tissue analysis [20e]. Following the same workflow, we first performed DDA analysis and then applied DDA-generated Hyp library to quantify Hyp substrates in tissues. The DDA-library based analysis only quantified 419 Hyp sites from all replicate analysis. Application of HypDB-generated spectral library increased the number of Hyp site quantifications by more than five times with 2510 sites (**Figure S8A**). Our result confirmed that HypDB-generated library greatly increased the Hyp sequence coverage and analysis sensitivity. DIA analysis with combined library generated by both HypDB and DDA analysis identified 2556 Hyp sites and 981 Hyp proteins (1% FDR). To test the reproducibility among replicate tissues, we performed correlation matrix analysis using the corrplot package in R. Our data showed that quantitative analysis of Hyp substrates allowed efficient clustering and segregation of tumor vs normal tissues (**Figure S8B**). After global q-value filtering and intensity normalization, we analyzed 1160 Hyp sites across all samples with pair-wise t-test and our analysis identified 12 upregulated sites and 24 downregulated Hyp sites in tumor (5% permutation-based FDR) (**Figure S8C and Table S5**). Functional enrichment analysis showed that Hyp proteins upregulated in kidney cancer were strongly enriched in cell-cell recognition and localized in axoneme (**Figure S8D**), while tumor suppressor ASPP1-SAM68 protein complex and p53-mediated signaling transduction processes were significantly enriched among down-regulated Hyp proteins in kidney cancer (**Figure S8E**).

## 3. Conclusions

A grand challenge in functional analysis of PTM pathways is the lack of annotation resources to profile modification substrates and annotate enzyme-target relationships. Proline hydroxylation is a key oxygen and metabolic-sensing posttranslational modification that governs the cellular programs in response to the hypoxia microenvironment and micronutrient stress. Earlier studies of proline hydroxylation mainly focused on its role in structural stability and maturation of cytoskeletal proteins such as collagens. In the past several decades, extensive biochemical studies on HIF pathways as well as other new Hyp substrates suggests that proline hydroxylation is widely involved in regulating protein-protein interaction, protein stability, signal transduction, metabolism and gene expressions. Growing evidence also suggested that specific Hyp pathways play critical roles in cancer development, metastasis, heart disease and diabetes. Systematic categorization and functional annotation of proline hydroxylation proteome will provide comprehensive understanding and important physiological insights into Hyp-regulated cellular pathways as well as potential therapeutic strategies targeting metabolic sensing Hyp pathways in diseases.

To address this need, we developed HypDB, an integrated online portal and publicly accessible server for functional analysis of proline hydroxylation substrates and their interaction networks. HypDB collected various data sources for comprehensive coverage of proline hydroxylation proteome, including manual curation of published literature, deep proteomics analysis of tissues and cell lines as well as integration with annotated Uniprot database. The site-localization and classification algorithm enabled efficient extraction of confident Hyp substrate identification from LCMS analysis. Our identification of highly confident Hyp substrates expanded the current annotation of human proline hydroxylation targets in Uniprot by over forty folds. Streamlined data processing and stoichiometry-based Hyp quantification allowed site-specific comparative analysis of Hyp abundance across 26 human organs and fluids as well as 6 human cell lines.

Bioinformatic analysis of the first draft of human proline hydroxylation proteome offer critical insights into the functional and structural diversity of the modification substrates. The analysis not only revealed diverse cellular pathways enriched with Hyp proteins including mRNA processing, metabolism, cell cycle and signaling, but also demonstrated for the first time that proline hydroxylation preferentially targets protein complexes and protein-interaction networks, indicating important roles of proline hydroxylation in fine-tuning protein structural features and mediating protein-protein interactions. Indeed, analysis of the expanded proline hydroxylation proteome with site-level secondary structure enrichment analysis indicated a significant enrichment of Hyp sites on alpha-helix, while site-level enrichment analysis of functional domains and regions revealed novel protein domain features that are preferentially targeted by proline hydroxylation, such as P-domain, NBD domain, ferritin-like domain and thioredoxin. These findings suggested potentially important roles for Hyp-mediated regulation of domain stability or activity that are worthy of further biochemical investigation.

MS-based analysis of Hyp proteome allows the stoichiometry-based quantification of Hyp abundance at the site-specific level. By classifying Hyp substrates based on stoichiometry dynamics, we revealed the enrichment of functional domains and activity with very high stoichiometry, indicating that proline hydroxylation on those domains may be required for the protein function, which is similar to collagen. In comparison, oxygen-sensing ODD domain was enriched with median stoichiometry and nucleotide or histone binding domains were enriched with low stoichiometry. Such difference may suggest differential activities of prolyl hydroxylases targeting various functional domains. Comparative analysis of Hyp stoichiometry across tissues also indicated variations in modification abundance at the site-specific level. Such variation may be attributed to the differential metabolic and gene expression profiles in various tissues.

The collection of MS-based identification of Hyp proteome in HypDB established annotated spectral library for Hyp-containing peptides that were identified and site-localized with high confidence. Such extensive spectral library enabled reliable and sensitive analysis of deep proteomic analysis of human cells and tissues with data independent acquisition. Application of the HypDB-generated spectral library in DIA analysis demonstrated excellent data reproducibility, significantly improved the coverage of Hyp proteome in cancer proteome analysis and revealed novel enrichment of Hyp sites that were significantly upregulated or downregulated in cancer tissues.

Although the current edition of HypDB is limited to human proteome, future development of HypDB will include proline hydroxylation proteome in other species. Comparative analysis of proline hydroxylation targets from diverse species will allow evolutionary conservation analysis of Hyp sites and identify functionally important Hyp targets in protein structure and activity. Further application of HypDB-generated spectral library in tissue analysis will enable the discovery of novel Hyp targets in disease animal model or patient samples and potentially lead to the development of clinically relevant theraupeutic strategies.

## 4. Experimental Methods

### 4.1 MS Raw data analysis

We collected MS data from the human proteome draft [23], deep proteome analysis of human cell lines [24], PHD interactome analysis [14, 25] and Hyp proteome analysis [8d] as well as IP-MS analysis of Flag-tagged HIF1A. All MS raw data collected above were searched with MaxQuant (version 1.5.3.12) against the Uniprot human database while having carbamidomethyl cystine as fixed modification and protein N-terminal acetylation, methionine oxidation, and proline hydroxylation as variable modification. Most of the raw data had trypsin as the digestion enzyme, while a few samples used other digestion enzymes, for example, LysC and GluC, based on the experimental procedure of original projects. Maximum missing cleavage number was set to 2 and the identification threshold was set at 1% false discovery rate for concatenated reversed decoy database search at protein, peptide and site levels.

### 4.2 Site localization classification and scoring

To filter out low confidence sites, we developed the site localization classification algorithm. Based on the experience that sites are localized more accurately when more ion fragments are found in corresponding MS2 spectra helping to localize the modification mass shift, our algorithm divided sites into three classes according to their modification localization confidence: exclusive localized sites in Class I, sites non-exclusive but distinguishable from similar modifications in Class II, and the rest in Class III. (**Figure 2A**)

For a site to be classified as Class I site, a pair of b-ions or y-ions separating the proline from other amino acids must be found to localize it exclusively. In this way, a mass shift caused by hydroxylation can only occur on that specific proline. And we gave credits to that ion pair in the scoring function for Class I sites as follows:

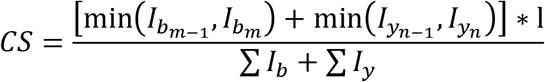

where *CS* stands for credit score, *I* stand for intensity of different ion fragments, for example, *I_b_m__* stands for the intensity of *b_m_*-ion, and *l* stands for peptide length. We gave credit to the pair of b-ions and y-ions that localize hydroxylation exclusively. The one with lower intensity within the pair will be selected, and we calculate the credit score based on the ratio of their intensities to average ion intensity on the same peptide.

Hydroxylation that cannot be exclusively localized but distinguishable from occurring on other prion-to-oxidize amino acid residuals are classified as Class II because we can infer that hydroxylation occurs on proline in this case. As all ions that separate proline from nearest amino acid may get oxidized easily, we gave credits to all ions that help to separate them in the scoring function for Class II sites as follows:

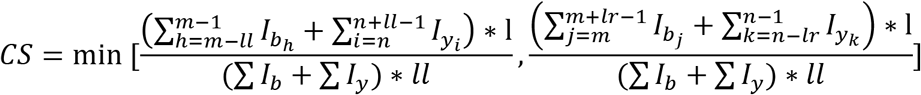

where *ll* and *lr* for distance between hydroxylated proline and nearest prion-to-oxidation amino acid residual on the left side and right side. Instead of only giving credit to the pair next to the side, for Class II sites, we gave credits to all ions that contributed to separate proline hydroxylation with other prion-to-oxidation amino acid residual. After that, we also calculate the ratio between the average intensity of selected ions and all ions on both sides, and the credit score is determined by the weaker side.

Sites that belong to neither Class I nor Class II are classified as Class III sites. There are chances that Class III sites are proline hydroxylation on other positions or other modifications that are identified falsely. Due to their low credibility, we do not score them and only use Class I and Class II sites for most of the following analyses.

### 4.3 Stoichiometry calculation

We calculate the stoichiometry of each hydroxyproline site according to the total peptide intensity and modified peptide intensity. For a specific site, we collect all modified and unmodified peptides that contain this site from MS data. Then, we get stoichiometries by dividing total modified peptide intensity by total peptide intensity. Site stoichiometries in different samples are calculated separately, so there might be multiple original stoichiometries for one site in the same tissue or cell line. We take the average stoichiometry for analysis in the following steps in this case.

### 4.4 Statistical enrichment of pathways, functional annotations, domains, and complexes

We use R packages including “GO.db”, “GOstats” and “org.Hs.eg.db” to perform enrichment analysis including Pfam, Kegg, and Gene Ontology – biological processes, molecular function and cellular compartment. We collected proteins of Class I and Class II sites from HypDB and performed a hypergeometric test for each term in the annotations above. Enrichment significance is log-transformed, and we used Benjamin-Hochberg correction to check the enrichment significance with a cutoff of 0.05.

Meanwhile, we performed enrichment tests by sample and stoichiometry quantiles respectively. For sample-specific enrichment tests, proteins with hyp sites discovered in different tissues and cell lines are analyzed respectively. While in the other group, we divide proteins into five quantiles according to the average stoichiometry of corresponding sites across all samples. Four cutoffs for quantiles are 5%, 30%, 70% and 95%. We also perform the log transformation and cluster the samples or quantiles according to the enrichment difference in different terms.

We also used Metascape for functional annotations and visualizations.

### 4.5 Motif enrichment analysis

The protein sequences of the proteins represented in HypDB were downloaded from the UniProt database. In-house Python scripts were written to extract peptides that contained Hyp sites that passed our stringent filtering criteria. These peptides were extended to the length of 13 or 27 amino acids and centered around the hydroxylated proline residue. The pre-aligned peptides were uploaded to the MoMo (Version 5.4.1) web application [18]. All protein sequences that were obtained from the UniProt database were set as the background for the analysis. Within the MoMo web application, the motif-x algorithm was selected. The minimum number of occurrences for a motif was set to 20. The sequence logos were generated by the MoMo web application.

### 4.6 Secondary structure analysis

The positions for the secondary structures of the proteins represented in HypDB were downloaded from the UniProt database. In-house Python scripts were developed to determine the number of Hyp sites and Non-Hyp sites found in secondary structure features for regions of proteins that have a known PDB structure.

### 4.7 Network connectivity analysis

All Class I, Class II, uniprot and literature sites in HypDB are collected and transformed into 7368 ENSP IDs with Uniprot. Then we look for interactions in the String database having having both nodes in the ENSP list, which is 16158 interactions respectively. To test the connectivity significance, we randomly picked 7368 ENSP ids from Uniprot proteins and counted interactions whose both nodes were included by the randomly selected sample in the String database. The pick and count process are repeated 10000 times, and these interaction counts from random samples are compared with the corresponding number of sites from HypDB.

We also built a protein-protein interaction network with these hyp proteins. From which we then selected some highly interconnected sub-networks which carry different biological functions with the help of Cytoscape software and the Mcode module.

### 4.8 Solvent accessibility analysis

With information from PDBe and Uniprot, we matched hydroxyproline proteins with corresponding pdb ID and protein structures in pdb files. Then, we use R package bio.PDB.DSSP to interpret pdb files that contain structural information and calculate the solvent accessibility of each Proline residual in the protein structure using the Sander and Rost accessible surface area (ASA) values. Then all accessibilities are divided by maximum accessibility of proline to get the relative accessibility number between 0 and 1.

### 4.9 Protein-protein interface analysis

The interacting domain pairs and instances of domain-domain interactions of three-dimensional protein structures were downloaded from 3DID (https://3did.irbbarcelona.org/index.php). In-house Python scripts were developed to analyze the number of Hyp sites interacting with another residue and the number of Hyp sites within three residues of an interacting residue.

### 4.10 Evolutionary conservation analysis

Evolutionary conservation analysis Hyp sites was performed using EggNOG ortholog database (v5.0) and EggNOG-mapper online portal [17]. Briefly, first, using EggNOG-mapper, Hyp proteins were mapped to the corresponding ortholog groups. Next, Hyp sites and non-Hyp proline sites on Hyp proteins were aligned to ortholog sequences using MAFFT algorithm [26]. The number of matches a Hyp site or non-Hyp proline site to a Proline for the same positions in ortholog sequences and the total number of sequences in the ortholog group were recorded. Lastly, HyperG test was performed for each Hyp site based on normalized number of matches to Proline residues in ortholog sequences for Hyp sites and non-Hyp sites, as well as the total number of any amino acid residues in ortholog sequences for the same position as the Hyp sites or non-Hyp sites.

### 4.11 Development of website and MySQL database

The website serves as a front-end interactive interface of the database. It was developed using HTML, CSS, Javascript, and PHP and works on a Linux-Apache-MySQL-PHP (LAMP) server architecture. The front-end was designed using the Bootstrap framework. Associated protein data are fetched using APIs from several sources. Protein sequences, identifiers, and descriptions are fetched from entries in the UniProtKB/Swiss-Prot knowledgebase [27] protein secondary structure data are fetched from PDBe [28] and domains are fetched from Pfam [29]. The protein sequences are displayed on the website using neXtProt Sequence Viewer (https://github.com/calipho-sib/sequence-viewer). The spectral graphs on the website are visualized using d3.js (https://d3js.org/). The backend of the website utilizes PHP to interface with a MySQL database that contains the data as shown in **Figure S2A**.

### 4.12 Transfection and Immunoprecipitation of HIF1A

Transfection and overexpression of Flag-tagged HIF1A was performed following a procedure as previously described [30]. Flag-tagged HIF1A plasmid (Sino Biological) was transfected into 293T cells with polyethylenimine. Cells were treated with 10 μM proteasome inhibitor MG-132 (Apexbio) for 4 hours prior to harvesting. Twenty-four hours after transfection, cells were washed with cold PBS buffer and lysed in lysis buffer (150 mM NaCl, 50 mM Tris-HCL, 0.5% NP-40, 10% glycerol, pH 7.5, protease inhibitor cocktail (Roche)) on ice for 15~20 mins. Then, the cell lysates were clarified by centrifugation prior to the incubation with anti-FLAG M2 affinity gel (Sigma) for 6 hours at 4 °C. After incubation, the M2 gel was washed with wash buffer (cell lysis buffer with 300 mM NaCl) for three times and then eluted with 3X Flag peptide (ApexBio). The eluate were mixed with 4 X SDS loading buffer and boiled, and then loaded onto homemade SDS-PAGE gel and stained with Coomassie blue (Thermofisher).

### 4.13 In-gel Digestion and LCMS analysis of HIF1A

A large gel piece covering a wide MW range above 100 kDa was cut out and subjected to reduction/alkylation and in-gel digestion with trypsin (Promega) as previously described [19b]. Tryptic peptides were desalted with homemade C18 StageTip and resuspended in HPLC Buffer A (0.1% formic acid) before being loaded onto a capillary column (75 μm ID and 20 cm in length) in-house packed with Luna C18 resin (5 μm, 100 Å, Phenomenex). The peptides were separated with a linear gradient of 7% to 35% HPLC Buffer B (0.1% formic acid in 90% acetonitrile) at a flow rate of 200 nl/min on Dionex Ultimate 3000 UPLC and electrosprayed into a high resolution Orbitrap Lumos mass spectrometer (Thermofisher). Peptide precursor ions were acquired in Orbitrap with a resolution of 120,000 at 200 m/z and peptides were fragmented with Electron Transfer/High Energy Collision Dissociation (EThcd) with calibrated charge-dependent ETD parameters and ETD Supplemental Activation, and acquired in Top12 data-dependent mode sort by highest charge state and lowest m/z as priority settings. Raw data was analyzed by Maxquant software following the same procedure and parameter setting as previously published dataset as described above.

### 4.14 Usage of HypDB website

A dedicated website with integrated MySQL database was established to host the HypDB service. The database schema includes four tables representing redundant Hyp site identifications, non-redundant Hyp site identifications, interaction interface analysis, evolutionary conservation analysis and solvent accessibility analysis. Each record in the site identification table is assigned a unique HypDB site ID. The website was designed with the Bootstrap framwork (v4.1.3) and features several key functions including a Search bar, Protein information page, Site information page, Database summary, Upload/contribute page and Download/export page.

Search bar allows the user to input an Uniprot accession number or Gene name of the protein of interest, and the server will use the information to extract and display a ranked list of most similar entries in real time. Clicking on an entry will bring the user to the protein information page where protein identifiers, description, and protein sequence are displayed. All Hyp sites are identified on the protein sequence as well as known acetylation and phosphorylation sites from PhosphoSitePlus database [31] are highlighted by different colors. The list of Hyp sites is further displayed below the sequence in the table which includes the site properties including localization class, localization score, stoichiometry, solvent accessibility and evolutionary conservation information. Hyp site table is followed by properties of Hyp proteins including protein-protein interaction, secondary structure, functional domains and domain-domain interactions. Hyp sites identified with MS/MS evidence in the HypDB have “Details” button displayed for each site in the site table on the protein information page. Clicking on the Details button will bring the user to the peptide information page where the best identified MS/MS spectrum for the site is displayed with annotations of fragment ions.

Contribute/Upload page allows the community to contribute raw MS/MS identifications to the HypDB through an embedded google form. Information regarding the raw data type, location, sample type, database searching parameters as well as user information will be entered into database. Raw data will be downloaded and processed using the same streamlined workflow. The data will pass through the classification and site-localization analysis process and annotated with the bioinformatic workflows as described above. The final data will be deposited into the HypDB to share with the research community.

Export/Download page allows the community to download the complete dataset deposited in the HypDB including both redundant and non-redundant modification site tables. In addition, the Export function will allow the user to select a list of proteins or tissues of interests and export the precursor ion m/z of selected Hyp proteins to set up targeted quantification method when acquiring data or export the collected spectral libraries of Hyp sites from the selected Hyp proteins to perform database searching with Data-Independent Acquisition (DIA).

### 4.15 Construction of DDA-based Spectral Libraries

To construct the study-specific DDA-based spectral libraries from the Kitata et al. and Guo et al. studies, a database search of the DDA data from each study was performed by MaxQuant (version 1.5.3.12). The parameters for the search engine were slightly modified from the parameters reported by the authors of each study. The maximum number of cleavages was set to two and the threshold for identification was set at 1% FDR. The variable modification of proline hydroxylation was included in addition to the variable modifications that the authors of each study reported. The spectral data for Hyp sites were compiled into an msp-formatted spectral library.

### 4.16 DIA Data Analysis

DIA data was analyzed using DIA-NN (v1.8) [32]. The default workflow for analysis using a spectral library was followed (https://github.com/vdemichev/diann). The DIA data from the Kitata et al. and Guo et al. studies were analyzed separately with DIA-NN. FDR (q-value) for protein groups and Hyp site identification was set at 1.0%. The analysis of the DIA data from each study was performed with spectral library from various source: HypDB Library, Study-Specific DDA-based Library and Combined Library generated by both HypDB and Study-Specific DDA Analysis. DIA-NN further applied global q-value filtering and intensity normalization to generate Hyp site matrix output for Hyp sites that were confidently quantified across all samples. Python scripts developed in-house to process the output from DIA-NN to be Hyp site-nonredundant. The matrix output from each study with nonredundant Hyp site quantification was used for clustering, annotation enrichment analysis and visualization using the Perseus software platform [33]. Missing values were imputed using a normal distribution, and the data was hierarchially clustered. The processed site-nonredundant Hyp intensity data from DIA-NN was also analyzed and visualized using R. Missing values were imputed using the K-Nearest Neighbor (KNN) method in the NAguideR tool [34].

## Supporting Information

Supporting Information is available from the Wiley Online Library or from the author. Supplemental figures include Figure S1. HypDB web portal with front page (top), protein-level view (bottom left) and peptide-level view (bottom right); Figure S2. Schema of the MySQL database of HypDB; Figure S3. Solvent accessibility and evolutionary conservation analysis of Hyp sites; Figure S4. Flanking sequence motif and neighboring protein modifications of Hyp sites; Figure S5. Examples of Hyp substrate proteins with site-specific Hyp stoichiometries in different tissues; Figure S6. Scatter plot showing the enrichment of CORUM protein complexes with Hyp substrate proteins; Figure S7. Hyp protein networks in enriched CORUM complexes; Figure S8. DIA analysis of Guo et al study of kidney cancer revealed differentially regulated Hyp substrates in tumor. Supplemental tables include Table S1. Current list of redundant Hyp sites collected in HypDB; Table S2. Current list of non-redundant Hyp sites collected in HypDB; Table S3. Site-specific stoichiometry distributions in tissues and cell lines; Table S4. Hyp identification and quantification from the DIA analysis of Kitata et al study of lung cancer tissues; Table S5. Hyp identification and quantification from the DIA analysis of Guo et al study of kidney cancer tissues

## Acknowledgements

We would like to thank the members of the Chen lab for helpful discussion and suggestion. We appreciate Christopher Lee of the College of Biological Sciences at the University of Minnesota for the help with the initial testing of the website. We would also like to thank Changzhi Wang for the initial development of evolutionary conservation analysis workflow in this study and Yi-Cheng Sin for helping with LCMS analysis. We are grateful for the Masonic cancer center and the Center for Mass spectrometry and Proteomics for providing LCMS access and analysis. This work was supported by the National Institute of Health (R35GM124896 to Y.C.).

Received: ((will be filled in by the editorial staff))

Revised: ((will be filled in by the editorial staff))

Published online: ((will be filled in by the editorial staff))

## Supporting Information

### Supplemental Information

#### Supplementary Figures

**Figure S1.**
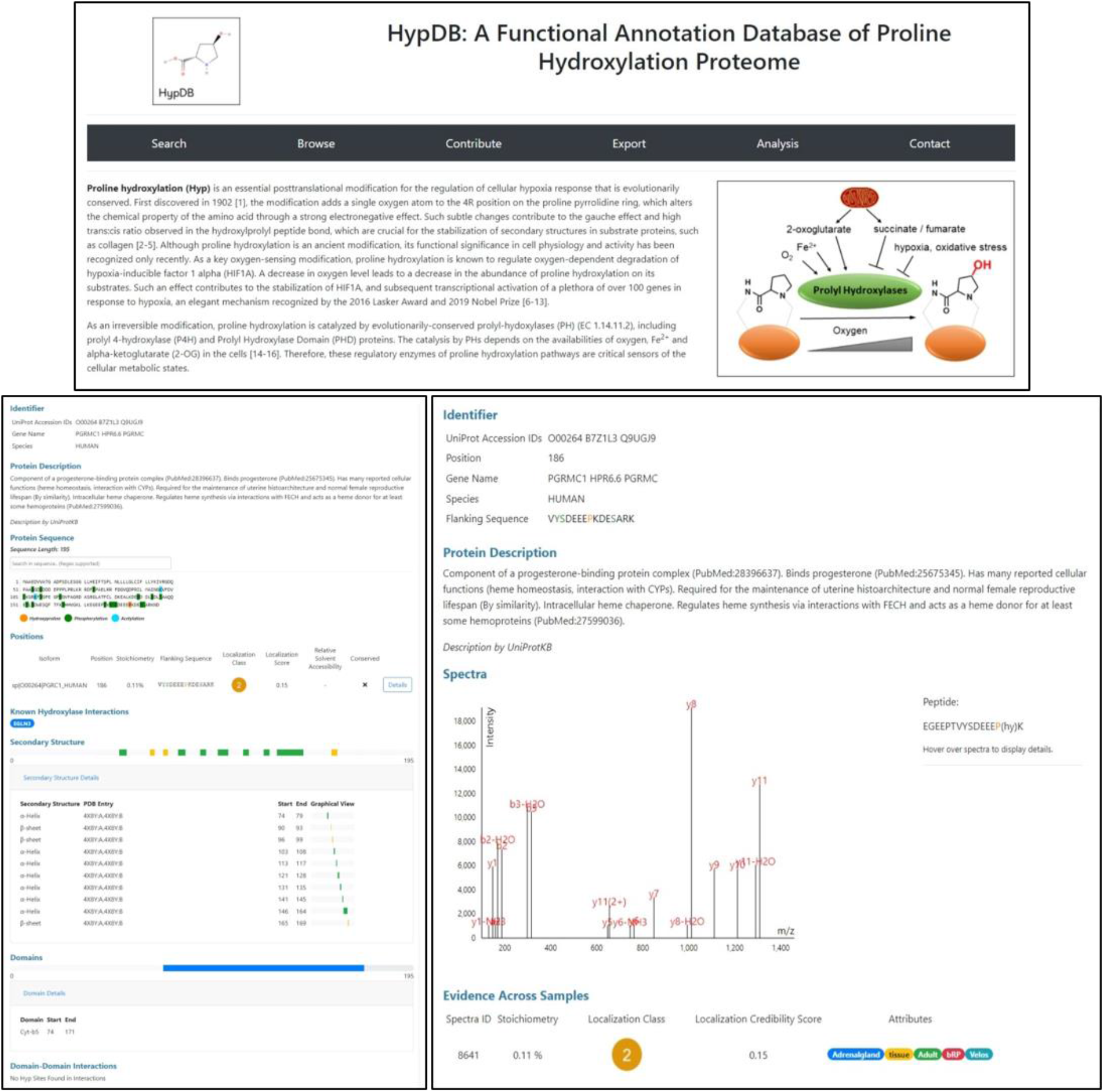
HypDB web portal with front page (top), protein-level view (bottom left) and peptide-level view (bottom right).

**Figure S2.**
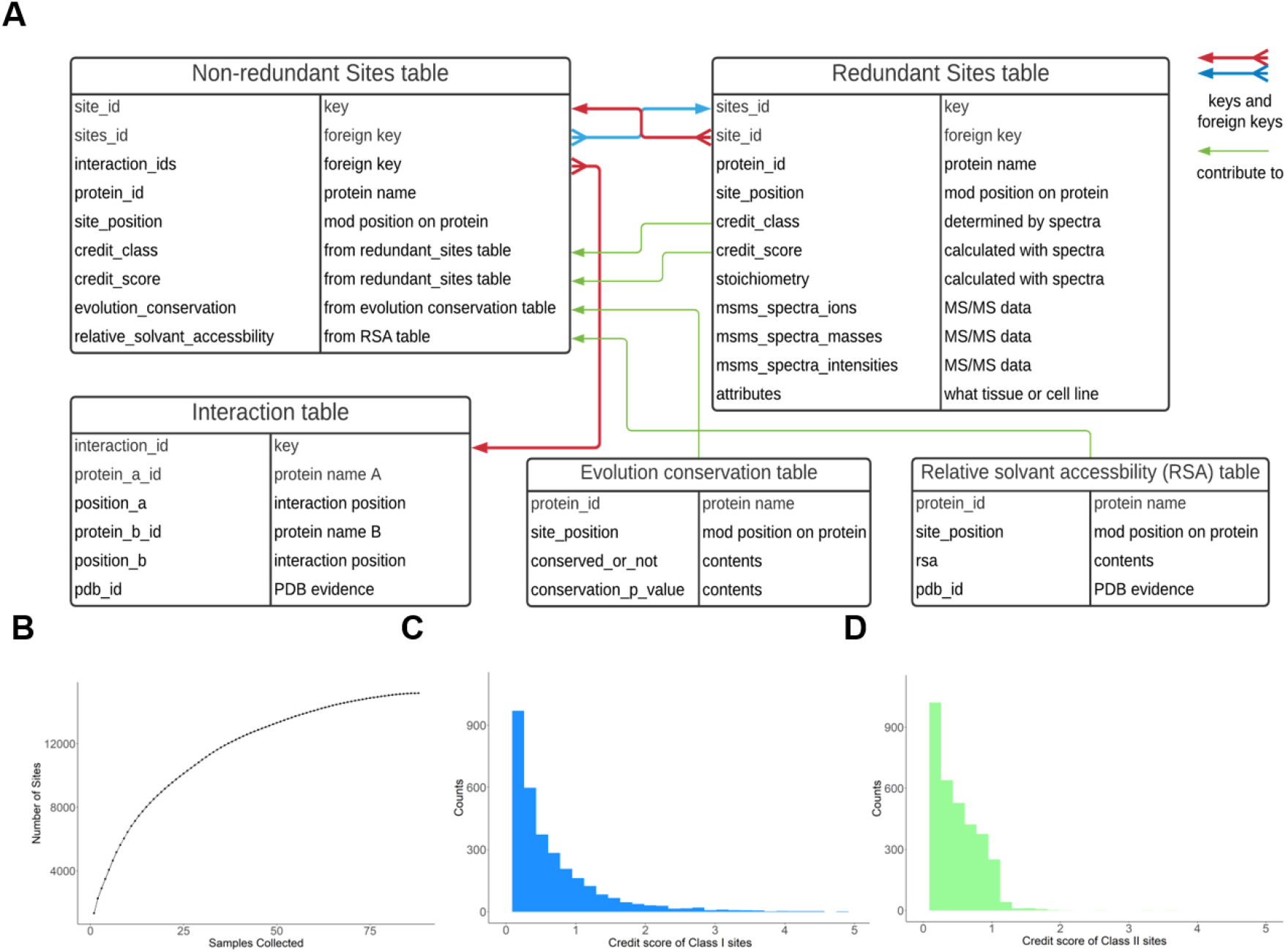
Schema of the MySQL database of HypDB and distribution of site properties. A. Simplified schema of tables in HypDB database; B. Accumulation of non-redundant Hyp sites as the number of samples collected from various tissues and cell lines increased; C. Distribution of the site localization credit score for Class I sites; D. Distribution of the site localization credit score for Class II sites.

**Figure S3.**
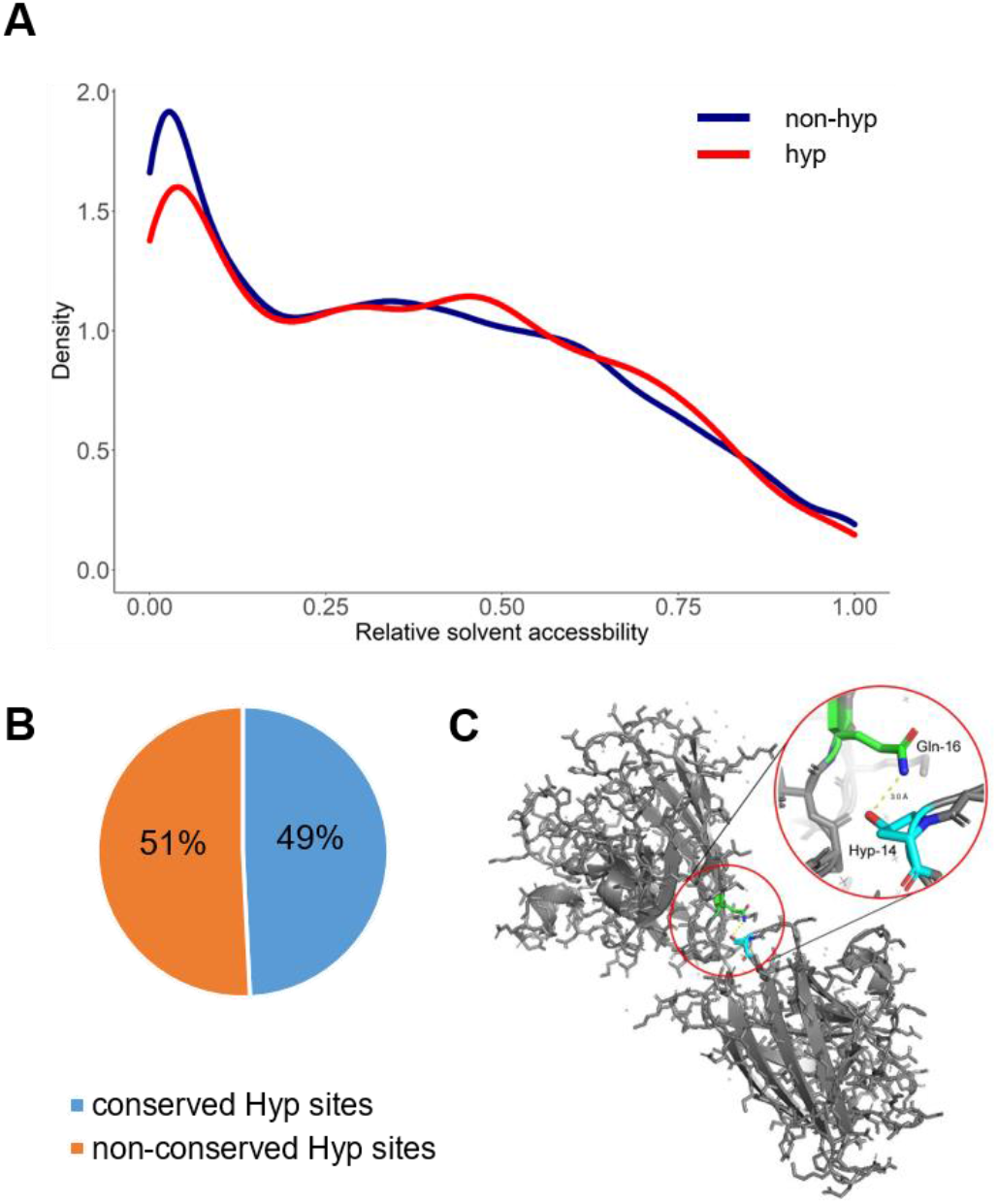
Solvent accessibility and evolutionary conservation analysis of Hyp sites. A. Relative solvent accessbiltiy distribution of hydroxyproline and non-hyp prolines on Hyp proteins; B. proportion of conserved and non-conserved hydroxyproline sites; C. Illustration of an interchain interaction between Gln-16 and Hyp-14 on SOD1.

**Figure S4.**
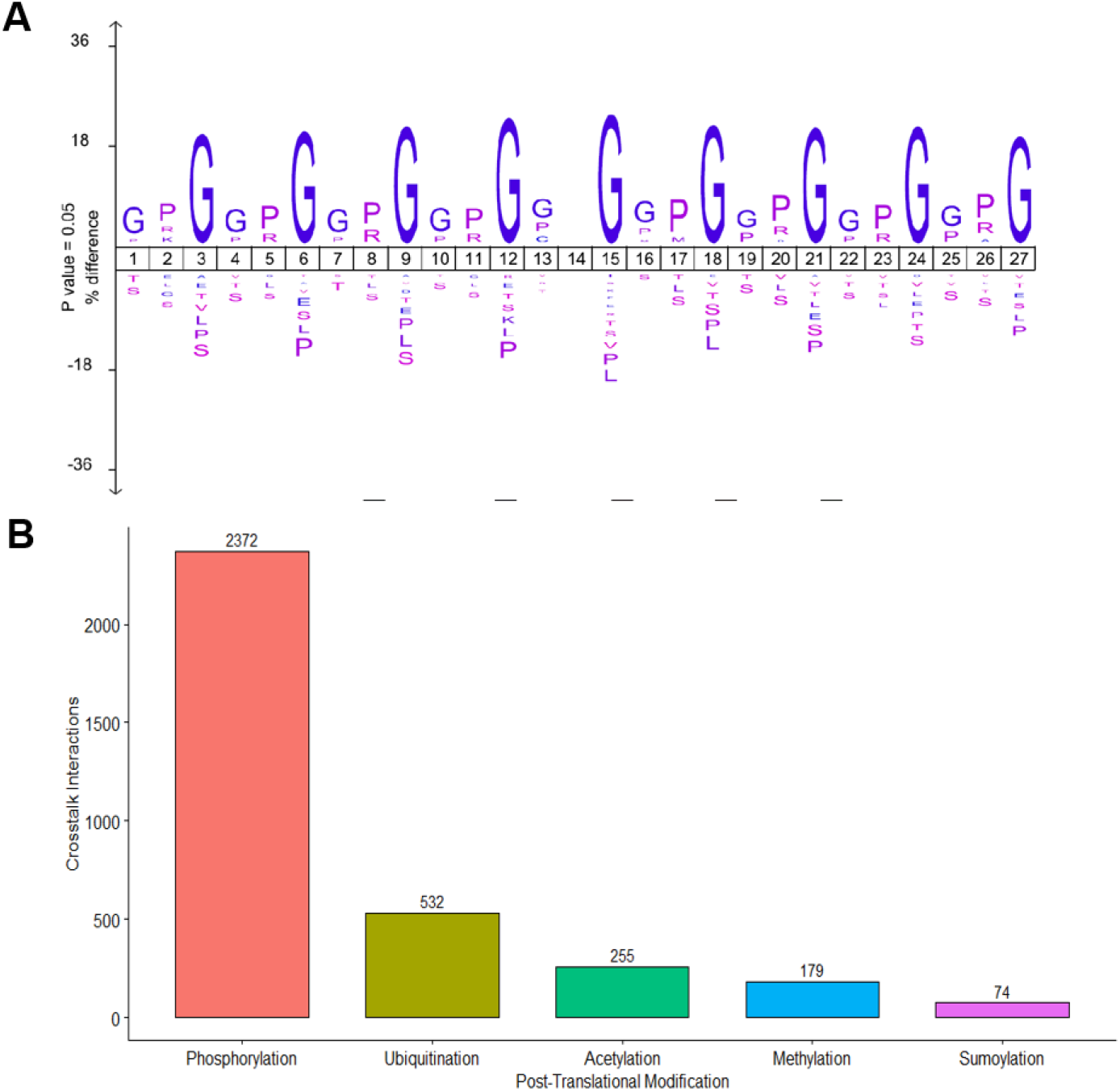
Flanking sequence motif and neighboring protein modifications of Hyp sites. A. Highly enriched PG motif with +/-− amino acids in Hyp proteome; B. Overlapping of Hyp sites with other PTMs sites in the neighboring positions of +/− 3 amino acids.

**Figure S5.**
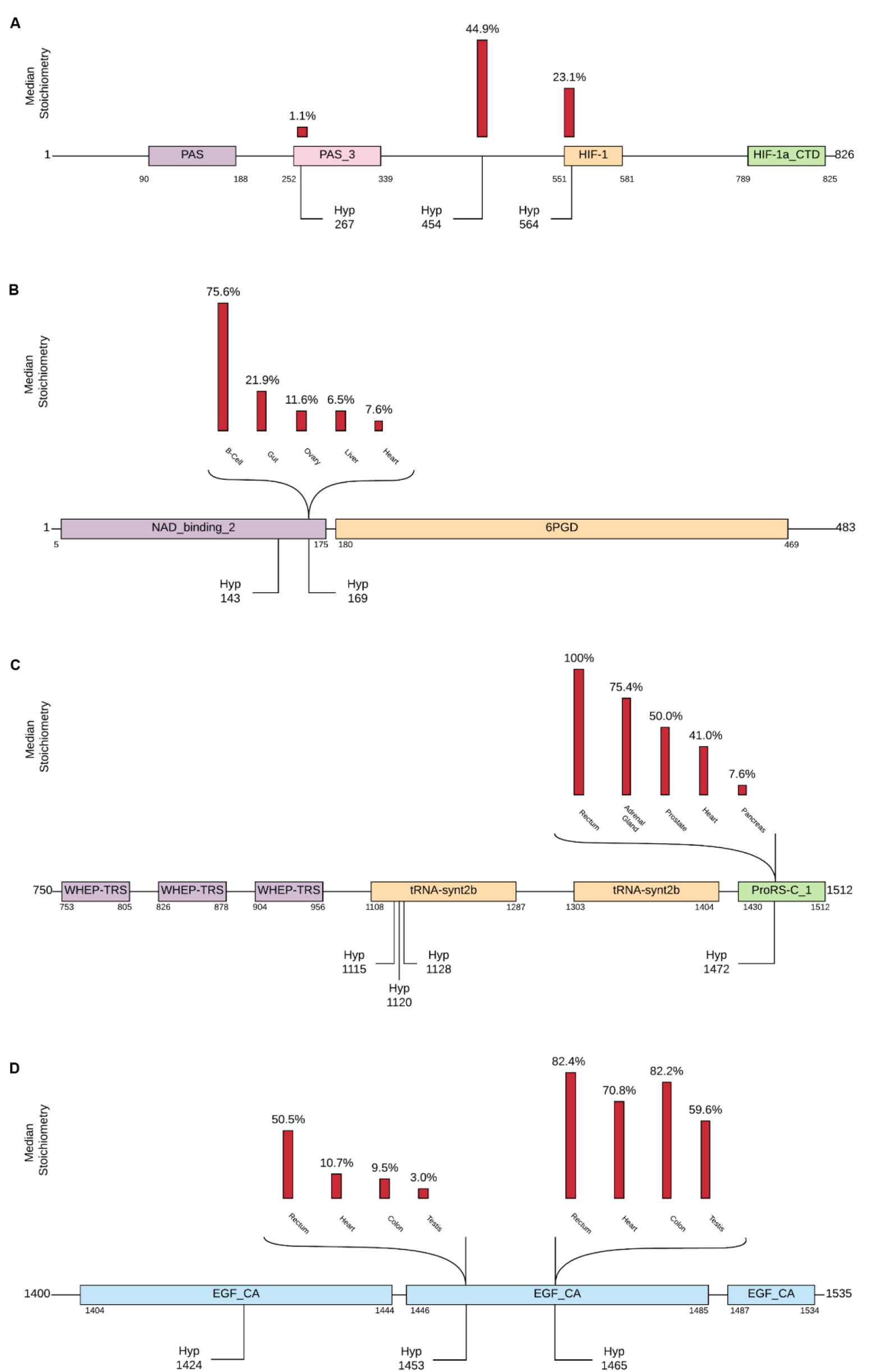
Examples of Hyp substrate proteins with site-specific Hyp stoichiometries in different tissues. A. Hypoxia-inducible factor 1A (Uniprot ID Q16665); B. 6-phosphogluconate dehydrogenase (Uniprot ID P52209); C. Bifunctional glutamate/proline-- tRNA ligase (Uniprot ID P07814); D. Fibrillin-1 (Uniprot ID P35555).

**Figure S6.**
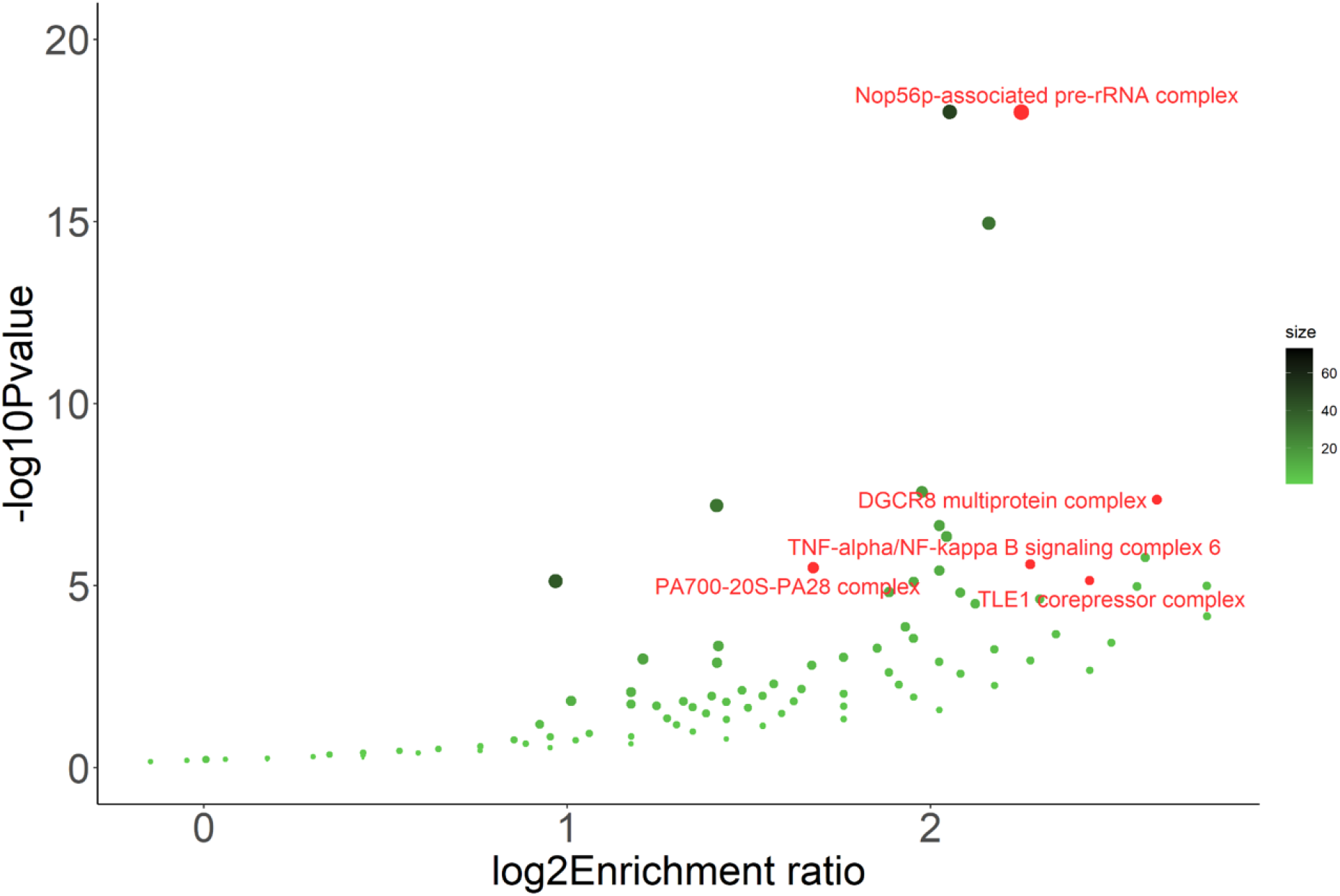
Scatter plot showing the enrichment of CORUM protein complexes with Hyp substrate proteins.

**Figure S7.**
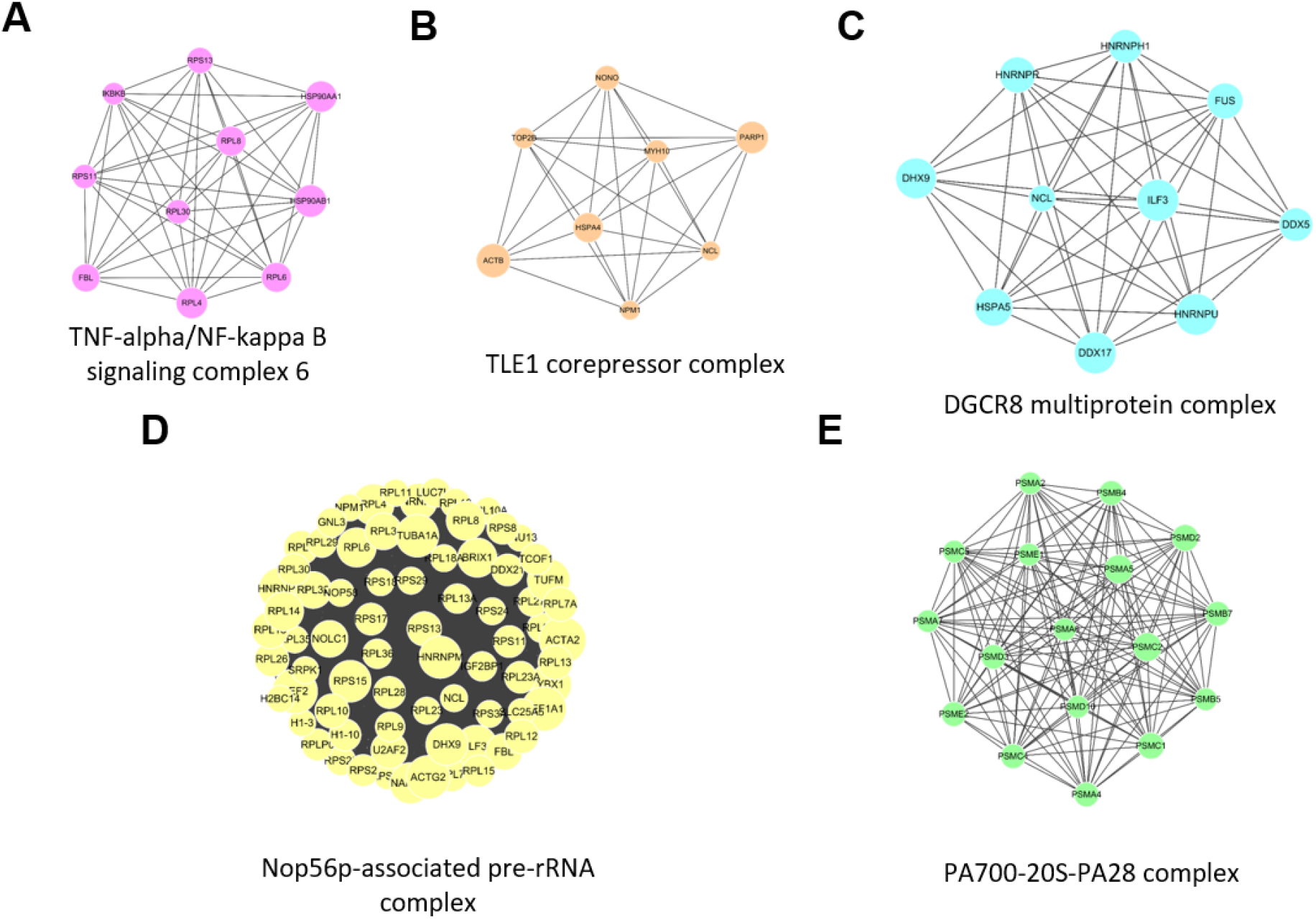
Hyp protein networks in enriched CORUM complexes. A. TNF-alpha/NF-kappa B signaling complex 6; B. TLE1 corepressor complex; C. DGCR8 multiprotein complex; D. Nop56p-associated pre-rRNA complex; E. PA700-20S-PA28 complex.

**Figure S8.**
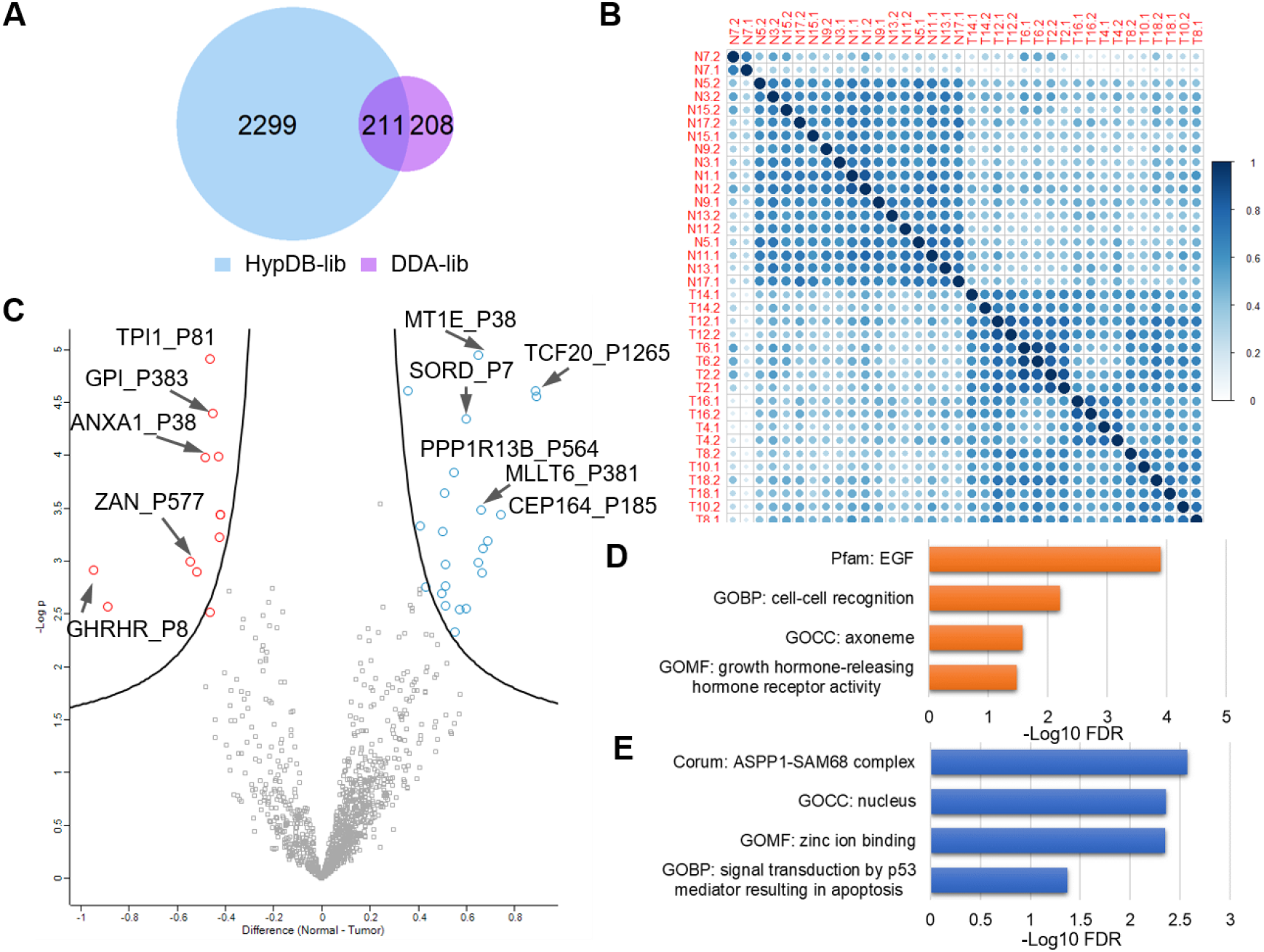
DIA analysis of Guo et al study of kidney cancer revealed differentially regulated Hyp substrates in tumor. A. Venn diagram of DIA identifiation of Hyp sites using HypDB-generated library and the library generated by the Data-dependent acquisition (DDA) of the study; B. Correlation plot showing clearly distinct classification of tumor and normal tissue analysis; C. Volcano plot of significantly up or down-regulated Hyp sites in normal and tumor tissues; Significantly enriched functional annotations among upregulated (D) and downregulated (E) proteins in tumor.

#### Supplementary Tables

**Table S1** Current list of redundant Hyp sites collected in HypDB.

**Table S2** Current list of non-redundant Hyp sites collected in HypDB.

**Table S3** Site-specific stoichiometry distributions in tissues and cell lines.

**Table S4** Hyp identification and quantification from the DIA analysis of Kitata et al study of lung cancer tissues.

**Table S5** Hyp identification and quantification from the DIA analysis of Guo et al study of kidney cancer tissues.

## Notes

### Competing Interest Statement

The authors have declared no competing interest.

